# Systematic functional genetic analysis of cell-substrate adhesion

**DOI:** 10.64898/2026.07.24.740569

**Authors:** Kaitlyn Manzer, Kuan-Chung Su, Matteo Di Bernardo, Heather R. Keys, Russell Walton, Brittania Moodie, Bingbing Yuan, Paul C. Blainey, Iain Cheeseman

## Abstract

Cellular adhesion is critical for tissue organization and integrity, but the full complement of proteins required for proper adhesion remains unresolved. Here, we define the requirements for cell-substrate adhesion in cultured human cells using orthogonal, large-scale functional genetic approaches. Using mechanical assays to test the maintenance (“shake-off”) or formation of adhesion and parallel large-scale assays of cell morphology, we identify dozens of gene targets with roles in adhesion. Our analyses reveal dynamic requirements for adhesion across timepoints and cell lines. We additionally conduct targeted downstream mechanistic analyses to resolve the molecular basis for altered adhesion. Collectively, we identify established and uncharacterized regulators of adhesion, including genes involved in mitosis, focal adhesions, actin regulation, and membrane trafficking. Unexpectedly, we find that cells that fail cytokinesis display impaired adhesion, with strongly altered actin organization and nuclear dynamics. Together, this work provides a comprehensive view of the genetic requirements for cell-substrate adhesion.

## Introduction

Cellular adhesion is an essential cellular process required for a wide range of physiological processes including wound healing, development, and cancer metastasis (Chastney et al., 2025). Cell-substrate adhesion is a highly regulated and dynamic process that requires cross talk from multiple structural and regulatory components. This includes focal adhesions – macromolecular transmembrane structures that link the extracellular matrix (ECM) to the intracellular cytoskeleton (Legerstee and Houtsmuller, 2021; Wehrle-Haller, 2012) (**Fig S1A)**. Focal adhesions are composed of integrins and multiple other proteins and connect to the actomyosin stress fibers that generate tension and control focal adhesion turnover (Bachmann et al., 2019; Hynes, 2002; Parsons et al., 2010). Adhesion is actively regulated to support different cellular processes. For example, during mitosis, cells undergo drastic remodeling of the focal adhesions and their cytoskeleton to round up, ensuring faithful chromosome segregation, while maintaining contact with the substrate through thin retraction fibers (Champion et al., 2017; Taubenberger et al., 2020) (**Fig S1A)**. Achieving a clear molecular understanding cell-substrate adhesion requires the comprehensive knowledge of the proteins that contribute to adhesion, both directly and indirectly. However, the molecular players that are required to ensure robust cellular adhesion remain incompletely defined.

Here, we develop large-scale functional genetics approaches to analyze cellular adhesion by leveraging the physical nature of adhesion in cultured cells. Pooled CRISPR/Cas9-based screens are powerful tools to interrogate gene function at scale (Shalem et al., 2014; Wang et al., 2014). However, traditional pooled CRISPR dropout screens evaluate relative cell fitness based on the loss of guide representation in cells cultured over a number of population doublings instead of interrogating cellular phenotypes. Our work establishes a robust “shake-off” strategy that enriches cells that are unable to maintain substrate adhesion under mechanical stress. In addition, we develop a complementary strategy that enriches cells that are unable to form initial attachments and adhesion following plating. Finally, we leverage optical pooled screening (OPS), which enables direct microscopy-based imaging to capture morphological information regarding cellular adhesion for each gene knockout. Our work reveals that gene knockouts that disrupt focal adhesions, proper mitotic progression, actin organization, and vesicle trafficking each result in reduced adhesion. Interestingly, we find that cytokinesis failure also results in the reduced maintenance of adhesion despite the absence of a mitotic arrest. Through downstream mechanistic analyses, we dissect the contributions of these pathways to cellular adhesion. Together, our work provides a comprehensive view of the molecular requirements for cell-substrate adhesion in cultured cells.

## Results and Discussion

### Pooled CRISPR screening identifies factors required for cellular adhesion

To identify genes required for cell-substrate adhesion in cultured cells, we designed a pooled CRISPR/Cas9-based strategy that takes advantage of the physical nature of adhesion (**Fig 1A**). For this analysis, we first used adherent, non-motile human cervical carcinoma HeLa cells expressing Cas9 under a doxycycline (DOX) inducible promoter (McKinley et al., 2015). We transduced cells with a pooled single guide RNA (sgRNA) library targeting all essential genes (Funk et al., 2022). At selected time points following Cas9 induction, we used mechanical agitation to detach weakly-adhered cells. By comparing sgRNA representation in the “shake-off” population to those that remained adhered to the plate (i.e., “shake-off” score), we identified multiple gene knockouts (KOs) that resulted in altered cellular adhesion (**Fig 1B, Table S1**). We observed a clear enrichment in the “shake-off” population for HeLa-expressed genes with established roles in cell-substrate adhesion (**Fig S1B**) (DepMap, 2025; Ly et al., 2024), such as the focal adhesion components Talin-1 (TLN1), Focal Adhesion Kinase (PTK2), and Integrin beta-5 (ITGB5). In contrast, genes with roles in cell-cell adhesion were not enriched in the shake-off screens, suggesting cell-cell interactions are dispensable for resisting mechanical detachment in this context.

**Figure 1:**
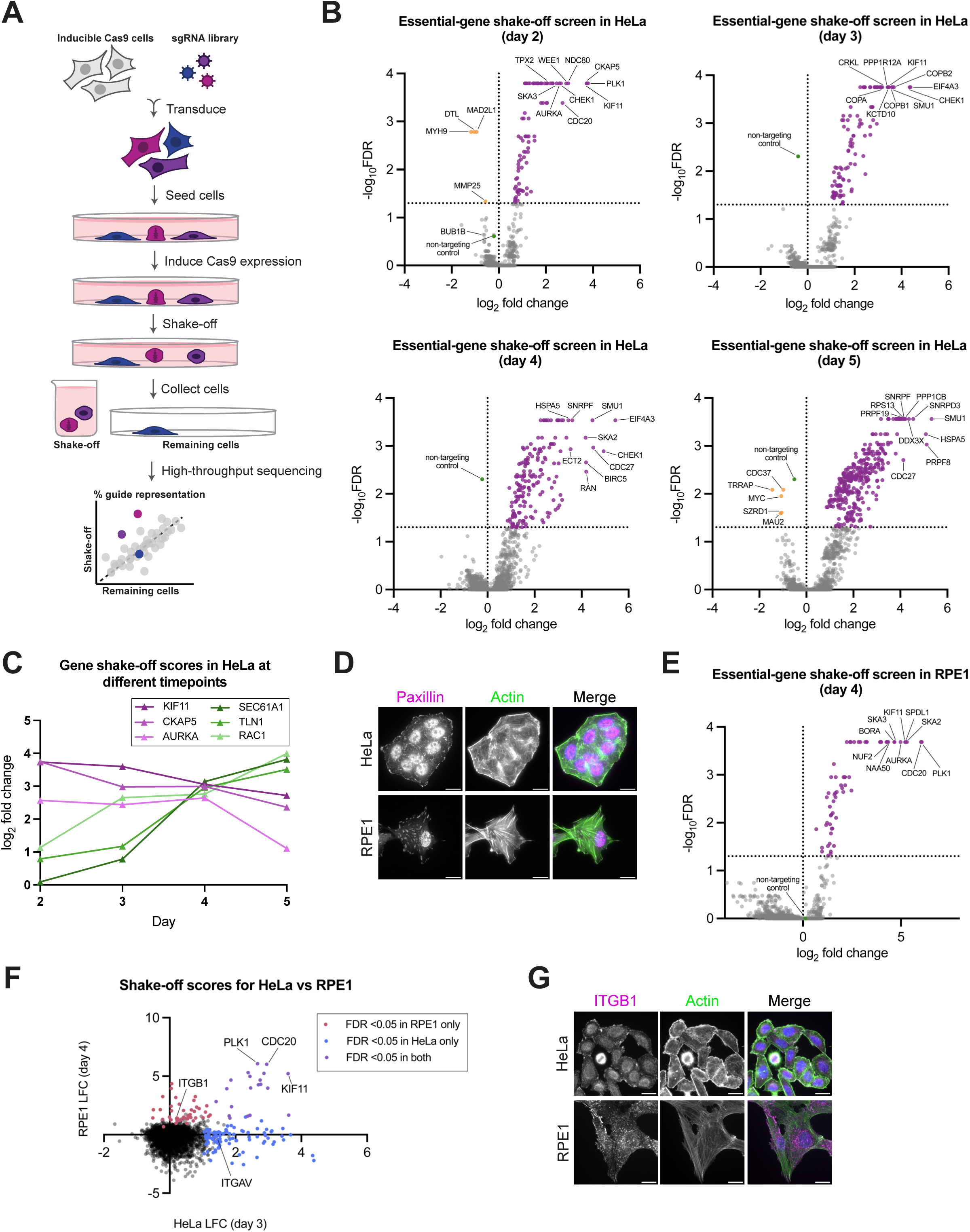
Shake-off CRISPR screening approach identifies factors required for cellular adhesion. a.) Schematic outlining shake-off screening approach. Inducible Cas9 cells were transduced with an sgRNA library. The cells were then seeded onto tissue culture dishes and treated with DOX to induce knockout for 2 to 5 days. Weakly adhered cells (e.g., mitotic cells or adhesion-defective cells, as depicted) were detached from the plate via shaking and washing. The detached cells (“shake-off”) and remaining cells (“remaining”) were each collected and sequenced to determine log_2_ fold change in guide representation between the shake-off and remaining populations. b.) Volcano plots of shake-off scores (LFC) for each targeted gene from the essential-gene shake-off screen in HeLa cells following a 2-, 3-, 4-, or 5-day induction time. False discovery rate (FDR) values were derived from MAGeCK enrichment analysis (positive LFC) or depletion analysis (negative LFC). A line is drawn at FDR = 0.05 to highlight the significantly enriched or depleted genes. Up to 10 significant genes (FDR < 0.05) with the strongest LFC are labeled in each direction. c.) Shake-off scores (LFC) for a subset of established mitotic (magenta) and trafficking or adhesion factors (green) at different time points in HeLa. d.) Representative z-projected immunofluorescence images of HeLa and RPE1 cells. Actin (phalloidin), paxillin (anti-paxillin). Merge also shows DNA (Hoechst) in blue. Scale bars, 20 µm. e.) Volcano plots of shake-off scores (LFC) for each targeted gene from the essential-gene shake-off screen in RPE1 cells following a 4-day induction time. FDR values were derived from MAGeCK enrichment analysis (positive LFC) or depletion analysis (negative LFC). A line is drawn at FDR = 0.05 to highlight the significantly enriched or depleted genes. Up to 10 significant genes (FDR < 0.05) with the strongest LFC are labeled in each direction. f.) Shake-off scores (LFC) from the essential-gene screen in HeLa cells (day 3) vs in RPE1 cells (day 4). Genes with a significant enrichment (MAGeCK FDR < 0.05) in each screen are highlighted. g.) Representative z-projected immunofluorescence images of HeLa and RPE1 cells. Actin (phalloidin), ITGB1 (anti-ITGB1). Merge also shows DNA (Hoechst) in blue. Scale bars, 20 µm.

Due to mitotic cell rounding and the reduced adhesion during mitosis, gene knockouts that extend mitotic duration are predicted to be enriched in the “shake-off” population. Indeed, we identified multiple genes whose depletion causes a mitotic arrest, including PLK1 and KIF11 (Blangy et al., 1995; Lane and Nigg, 1996) (**Fig 1B**). Conversely, knockouts of the spindle assembly checkpoint genes MAD2L1 and BUB1B, which shorten mitotic duration (Gorbsky et al., 1998; Meraldi et al., 2004), were depleted from the “shake-off” population at the earliest time point (**Fig 1B**). Thus, this large-scale approach is effective in identifying genes whose loss alters adhesion both directly and indirectly. Gene set enrichment analysis (GSEA) of gene knockouts with reduced adhesion further revealed pathways related to the actin cytoskeleton, vesicle trafficking, protein translation, and mRNA splicing (**Table S2**).

We next sought to test the dynamic changes that occur to cellular adhesion following the induction of the gene knockouts. The time required for phenotypes to manifest varies across gene targets due to the kinetics of protein depletion, the degree of protein depletion needed for a phenotype to be observed, and both the primary and secondary consequences of the knockout on cellular function. Based on the analysis of different time points following Cas9 induction, we uncovered additional modulators of cellular adhesion (**Fig 1B**). This temporal analysis additionally revealed differences in the kinetics of the shake-off scores for different knockouts. We observed a general trend in which mitotic genes were enriched in the shake-off at earlier time points but then became depleted from the population (**Fig 1C**), suggesting the loss of these knockout cells due to lethality. Conversely, established players in cellular adhesion and vesicle trafficking tended to become enriched in the shake-off population at later time points (**Fig 1C**), suggesting the need to substantially deplete the existing protein pool. Together, these results demonstrate the robust ability of this mechanical enrichment strategy to identify gene targets whose loss dynamically alters cell-substrate adhesion.

### Shake-off analysis uncovers cell-type specific adhesion requirements

Adhesion is a highly-regulated process that varies across cell types to support their unique functional requirements and cell-substrate interactions. To assess these behaviors in a different cell line, we next tested non-transformed retinal pigment epithelial RPE1-hTERT cells (Bodnar et al., 1998), which exhibit cell motility, are strongly adherent, and display distinct actin cytoskeleton and focal adhesion organization relative to HeLa cells (**Fig 1D**). We performed a shake-off analysis in RPE1 cells using the essential-gene library (**Fig 1E, Table S1**). We selected a time point following 4 days of Cas9 induction to match the extent of protein depletion at day 3 in HeLa cells, a time point that had showed clear enrichment in the shake-off with minimal gene depletion (**Fig S1C**) (McKinley and Cheeseman, 2017). We confirmed the similarities between these time points based on the degree of sgRNA depletion (**Fig S1D**).

By comparing the results from the shake-off analysis in HeLa and RPE1 cells, we observed multiple differences in cell-line specific adhesion. First, the total number of cells that detached with mechanical shake-off was significantly reduced in RPE1 cells, reflecting their increased adherence relative to HeLa cells. This limited our ability to recover sufficient coverage of “shake-off” cells over the guide library (∼50-fold), and therefore we have been cautious in making conclusions regarding the absence of a phenotype in RPE1 cells. Second, based on the analysis of genes enriched in the “shake-off” population in RPE1 cells, we observed that nearly every hit was an established mitotic factor, with a majority of terms enriched in the GSEA pertaining to cell division (**Table S2**). In contrast, many of the other gene targets that we identified in the HeLa screen, such as membrane trafficking genes, were absent. We speculate that this may be due to an increased redundancy in the adhesion machinery or the strongly adherent nature of RPE1 cells. Third, we identified differential gene requirements between HeLa and RPE1 cells. In particular, the integrin subunit integrin beta-1 (ITGB1) was enriched in the RPE1 screen, but not in the HeLa analysis at any time point (**Fig 1F, Fig S1E**). Consistent with this, immunofluorescence analysis showed that HeLa cells possess fewer ITGB1-containing focal adhesions than RPE1 cells (**Fig 1G**). Conversely, integrin alpha-V (ITGAV) was enriched in the HeLa screen, but not in the RPE1 screen (**Fig 1F**). This suggests that RPE1 and HeLa cell lines have unique requirements for integrin subunits for substrate adhesion. In contrast, the majority of the other hits in the RPE1 shake-off analysis were also identified in the HeLa shake-off screens at least at one time point (**Table S1**). These results provide insights into how adhesion is regulated across cell lines and demonstrate the utility of employing this mechanical enrichment strategy to study adhesion across different cellular contexts.

### Optical pooled screening identifies mitotic-defective shake-off hits

To understand the mechanisms underlying the adhesion defects for the gene targets identified in our shake-off analysis, we next leveraged orthogonal large-scale microscopy-based analyses. We first utilized a published optical pooled screening (OPS) dataset from our recent work (Funk et al., 2022), referred to here as ‘Vesuvius’. The Vesuvius dataset used the same HeLa cell line and essential-gene sgRNA library, but measured microscopy-based morphological features following 78-hours of Cas9 induction, enabling direct comparison with our 3-day (72-hour) shake-off analysis (**Fig 2A**). We first sought to distinguish gene knockouts whose enrichment in the shake-off population can be explained by prolonged mitosis. Vesuvius used a classifier trained on DNA features to determine the percentage of cells in mitosis for a given knockout, generating a “mitotic index” measurement for each gene. We observed a clear relationship between an increased mitotic index and shake-off enrichment, particularly for established players in mitosis (**Fig 2B**). This comparative analysis also demonstrated the advantage of orthogonal strategies to evaluate mitotic enrichment. For example, despite having apparently normal mitotic indices in Vesuvius, we observed an enrichment for the cell cycle regulators WEE1 and CHEK1 in the shake-off analysis (**Fig 2B**). We further observed a clear mitotic arrest in individual knockout cells based on a rounded cell morphology (**Fig S2A**). In this case, we found that the presence of uncondensed DNA during mitosis likely caused these knockouts to evade the mitotic classifier and prevent their detection in the imaging-based morphological analysis (**Fig S2B**). For WEE1, this mitotic chromosome morphology has been attributed to chromosome pulverization driven by replication stress and DNA damage (Haykal et al., 2024), and our analysis further extends this role to CHEK1. In total, based on the orthogonal mitotic index analysis, we find that ∼38% of the day-3 shake-off hits are likely to display reduced adhesion due to an increased mitotic duration.

**Figure 2:**
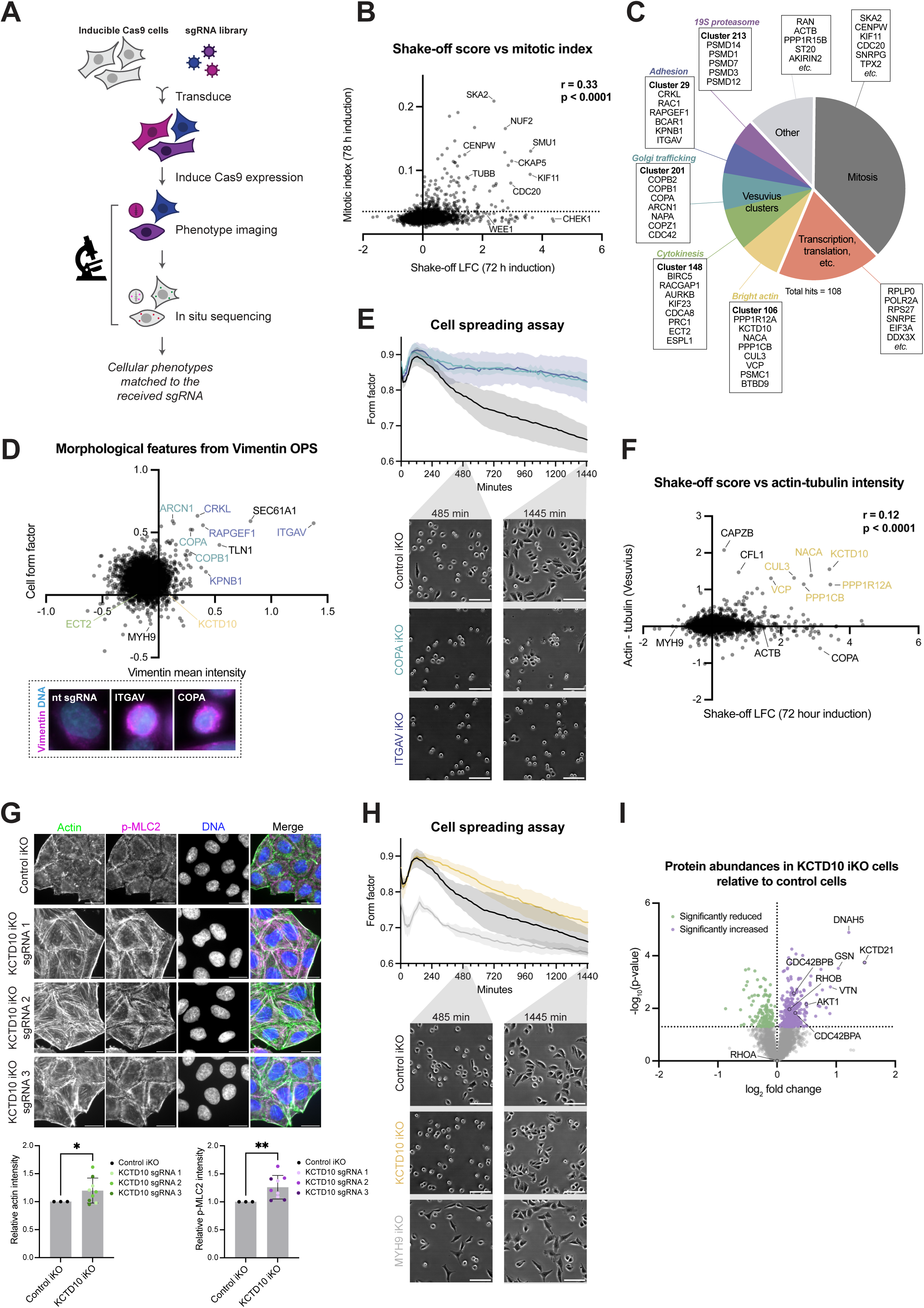
Shake-off hits have underlying defects in mitosis, gene expression, adhesion, trafficking, actin, and cytokinesis. a.) Schematic outlining optical pooled screening approach. Inducible Cas9 cells were transduced with an sgRNA library. Following a 78-hour treatment with DOX to induce knockout, cells were fixed and imaged, first for morphological phenotyping and then for in situ sequencing to determine sgRNA identity. b.) Shake-off scores (LFC) in the essential-gene shake-off screen in HeLa cells (72-hour induction) vs mitotic index from the essential-gene optical pooled screen published on https://vesuvius.wi.mit.edu/ (78-hour induction). Pearson correlation. Line at 0.0354 marks the threshold used to classify knockouts as having an elevated mitotic index in the following analyses. c.) Pie chart categorizing the 108 significantly enriched (LFC > 0, FDR < 0.05) knockouts in the essential-gene shake-off screen in HeLa cells at the 3-day time point. Genes with elevated mitotic indexes (≥0.0354) in Vesuvius that are likely enriched due to mitotic rounding were classified as “Mitosis”. Remaining hits with known roles in core cellular processes (transcription, translation, splicing, DNA replication, and protein folding) were grouped as “Transcription, translation, etc.”. Remaining hits were categorized by their corresponding phenotypic clusters in Vesuvius, with highly represented clusters (>2 genes) labeled, and others assigned as “other”. Phenotypic clusters are labeled by the shared pathway or phenotype. d.) Mean vimentin intensity versus cell form factor from vimentin optical pooled screen. Select genes are labeled and colored by phenotypic clusters as in Fig 2C. Representative images of cells that received nontargeting guides or guides targeting ITGAV or COPA are shown. e.) Cell spreading assay with control, ITGAV, and COPA iKO HeLa cells. Spreading assays with iKO cells were performed in parallel; the same control data are shown in each figure panel for comparison. (Top) Quantification of cell form factor of cells over time following trypsinization and re-plating onto fibronectin-coated surfaces, averaged across three experimental replicates (*n* > 150 per condition per replicate). Data represent mean ± SEM. (Bottom) Representative brightfield images of cells at select time points. Scale bars, 100 µm. f.) Shake-off scores (LFC) in the essential-gene shake-off screen in HeLa cells (72-hour induction) vs the mean actin intensity minus mean tubulin intensity from the essential-gene optical pooled screen published on https://vesuvius.wi.mit.edu/ (78-hour induction). Pearson correlation. g.) (Top) Representative z-projected immunofluorescence images of control and KCTD10 iKO HeLa cells following 72 hours induction. Actin (phalloidin), phospho-myosin light chain 2 (anti-p-MLC2), and DNA (Hoechst). Scale bars, 20 µm. (Bottom) Quantification of the relative mean p-MLC2 and actin immunofluorescence intensity in control and KCTD10 iKO cells across three experimental replicates (*n* > 200 per condition per replicate). Within each replicate, values are normalized to the average control value. Error bars represent standard deviation. Two-tailed Welch’s t-tests were performed (ns, not significant; *p ≤ 0.05, **p ≤ 0.01, ***p ≤ 0.001; ****p ≤ 0.0001). h.) Cell spreading assay with control, KCTD10, and MYH9 iKO HeLa cells. Spreading assays with iKO cells were performed in parallel; the same control data are shown in each figure panel for comparison. (Top) Quantification of cell form factor of cells over time following trypsinization and re-plating onto fibronectin-coated surfaces, averaged across three experimental replicates (*n* > 150 per condition per replicate). Data represent mean ± SEM. (Bottom) Representative brightfield images of cells at select time points. Scale bars, 100 µm. i.) Volcano plot comparing protein abundance in control and KCTD10 iKO HeLa cells as measured by quantitative TMT-MS. Proteins with significant changes in abundance as determined by a Student’s t-test (p < 0.05) are highlighted. Select proteins are labeled.

### Adhesion-defective knockouts disrupt actin organization, vesicle trafficking, and cytokinesis

We next sought to define the mechanisms beyond a mitotic arrest that underlie the remaining adhesion defects identified in our large-scale analysis. Many of these gene knockouts target core cellular processes, such as transcription, translation, protein folding, and splicing, and likely impair adhesion as an indirect downstream consequence of altered protein production (**Fig 2C**). To investigate the remaining 47 hits, we leveraged the phenotypic clustering from Vesuvius, which groups gene targets based on their phenotypic similarity to reveal functional relationships (Funk et al., 2022). Interestingly, the majority of our shake-off hits corresponded to just a few phenotypic clusters, likely representing different pathways required for cellular adhesion. One highly-represented cluster grouped established direct adhesion factors, such as ITGAV and the small GTPase Rac1 (**Fig 2C**). The other clusters group genes involved in Golgi trafficking, cytokinesis, the 19S proteasome, or whose knockouts result in enhanced F-actin staining (**Fig 2C**). Notably, recent work from our group demonstrated that the loss of the 19S proteasomal lid subunits causes monopolar mitotic spindles and a mitotic arrest (Marescal and Cheeseman, 2025), likely explaining the decreased adhesion.

To determine how disruptions to these genes result in adhesion defects, we conducted a new optical pooled screen designed to assess cell morphology and adhesion with our essential-gene library using vimentin staining to mark the cell body (**Fig 2D**, **Table S3**). Our quantitative analysis of morphological parameters revealed that the loss of established adhesion factors and focal adhesion proteins resulted in cells that are smaller, rounder, and subsequently brighter, consistent with reduced cell spreading (**Fig 2D**). These gene targets formed a coherent gene cluster based on their similar morphological phenotypes [AXIN1; C16orf86; CEBPB; CHD4; CRKL; DDX17; DDX3X; ERP44; FERMT2; ILK; ITGAV; ITGB1; ITGB5; KLHL22; MEN1; NCBP1; NCBP2; NFE2L2; RAC1; RAPGEF1; RSU1; SAP130; SEC61A1; SEC61G; SEC62; SRP54; SUCO; TLN1; TMED10; TNS], supporting their shared roles (**Table S3**). We propose that this cluster represents the core module required for cell-substrate adhesion. Notably, this grouping based on cell morphology also contains components of the secretory pathway. In contrast, other hits identified in our shake-off screen, such as those involved in actin organization or cytokinesis, did not display a similar cell morphology.

To assess the role of vesicle trafficking in cellular adhesion, we utilized a cell spreading assay in which we plated trypsinized knockout cells on a fibronectin-coated surface and evaluated their ability to reform adhesions and spread out. We quantified the degree of cellular spreading by measuring cell roundness using the form factor in CellProfiler (Stirling et al., 2021), in which perfect circular objects will have a form factor of 1, and more elongated or irregular objects have a form factor of less than 1. As expected, knockout cells for the integrin subunit ITGAV, which has an established role in adhesion (Hynes, 2002), exhibited an impaired ability to attach and spread out in this assay (**Fig 2E**). Similarly, knockouts for the COPI subunit COPA, which is involved in retrograde and intra-Golgi trafficking, also displayed a reduced ability to spread (**Fig 2E**), supporting its role in adhesion dynamics. COPA knockout cells do not result in a mitotic arrest based on immunofluorescence analysis (**Fig S2C**), but do exhibit an elongated morphology, likely reflecting a loss of adhesion at the cell surface. We hypothesize that reduced adhesion and altered cell morphology reflect a consequence of impaired trafficking and secretion of cellular adhesion molecules, such as integrins or ECM components, to the cell surface, causing these knockouts to phenocopy the loss of adhesion components. Overall, these dual mechanical and microcopy-based approaches identified both established and unexpected contributors to cell adhesion, underscoring the power of utilizing orthogonal approaches to study cellular phenotypes.

### Dysregulation of actin and contractility impairs cellular adhesion

Focal adhesions require an interaction between integrins and the actin cytoskeleton, with stress fibers acting to stabilize mature focal adhesions (Bachmann et al., 2019; Hynes, 2002). Consistent with a role of the actin cytoskeleton in promoting cellular adhesion, we found that actin (ACTB) was required for adhesion in our shake-off analysis (**Fig 2C**). To investigate the interplay between actin organization and cellular adhesion, we evaluated filamentous actin (F-actin) levels from the Vesuvius data set relative to the mechanical shake-off scores. Interestingly, we found that there was a clear relationship between actin levels and adhesion, with increased actin staining relative to tubulin levels being associated with reduced cellular adhesion (**Fig 2F**). Although this may be counterintuitive given the established role of stress fibers in cell-substrate anchoring, increased actin assembly or stability may impact adhesion by altering actin dynamics and disrupting adhesion mechanics.

Amongst these gene targets, we found that KCTD10 knockout cells exhibited the brightest actin staining in the Vesuvius dataset and displayed significant enrichment in the shake-off screen. KCTD10 acts as a substrate adaptor for CUL3 in a Cullin-RING E3 ubiquitin ligase (CRL), which targets substrates for degradation through the 26S proteasome. To assess the role of KCTD10 in adhesion, we first imaged individual inducible knockout (iKO) cells, which revealed increased actin stress fiber formation (**Fig 2G**) and elevated levels of active myosin II based on staining for phosphorylated myosin light chain 2 (p-MLC2) (**Fig 2G**). These findings suggest that KCTD10 cells display enhanced stress fiber formation and increased contractility. Other shake-off hits in the KCTD10-containing Vesuvius cluster include players in CUL3-mediated protein degradation, including CUL3 itself and the substrate adaptor BTBD9 (Freeman et al., 2013; Genschik et al., 2013), as well as the regulatory and catalytic subunits of myosin phosphatase, PPP1R12A (MYPT1) and PPP1CB (Kiss et al., 2019). This suggests that loss of these genes results in excessive myosin phosphorylation and increased actin stress fiber formation, resulting in impaired adhesion. Myosin II proteins are actin-based motor proteins with established roles in generating contractile forces (Murrell et al., 2015). We reciprocally found that the myosin IIA subunit MYH9 was depleted in our shake-off analysis (**Fig 1B**), possibly reflecting an enhancement in adhesion.

To evaluate the contribution of MYH9 and KCTD10 to cellular adhesion, we used the cell spreading assay described above. We observed a reduced ability of KCTD10 iKOs to spread, supporting their altered adhesion mechanics (**Fig 2H, Fig S2D**). Conversely, MYH9 iKOs displayed much faster spreading, consistent with previous reports (Betapudi, 2010) (**Fig 2H**). Importantly, neither of these knockouts exhibited a mitotic arrest (**Fig S2C**). Together, these results highlight the importance of actin regulation and actomyosin contraction in cellular adhesion.

To identify potential KCTD10 substrates that could regulate actin organization, we performed whole-cell proteomic analysis of control and KCTD10 knockout cells (**Fig 2I**, **Table S4**). Based on quantitative mass spectrometry, we identified multiple proteins with increased abundance in the KCTD10 iKO cells that could represent KCTD10 substrates that contribute to these phenotypes. These include the CDC42 effectors CDC42BPA (MRCK-α) and CDC42BPB (MRCK-β), which phosphorylate myosin light chain (Unbekandt and Olson, 2014), as well as AKT1, which has roles in adhesion regulation (Higuchi et al., 2013; Somanath et al., 2007; Wang and Basson, 2011). We also detected a small but significant increase in RhoB levels, consistent with prior work suggesting that KCTD10 regulates RhoB in contexts such as endothelial barrier function, developmental cell fusion, and breast cancer (Kovacevic et al., 2018; Murakami et al., 2019; Rodríguez-Pérez et al., 2021). Interestingly, one of the proteins with the strongest increase in abundance was KCTD21, a related CUL3 adaptor protein. This may reflect KCTD10-mediated degradation of KCTD21, similar to what has been described for KCTD13 (Cheng et al., 2024). Overall, these studies implicate KCTD10 as a major regulator of actin organization and contractility across broad cellular contexts, with important roles in maintaining cellular adhesion.

### Genome-wide analyses distinguish regulators of initial cell attachment versus adhesion maintenance

The essential-gene library used for the large-scale analyses described above represents a set of genes whose loss has consequences on cellular viability (Funk et al., 2022). To identify additional gene targets that alter cellular adhesion without resulting in strong fitness defects in cell culture, we next sought to employ the shake-off strategy to evaluate the contribution of every human gene to cell-substrate adhesion. We conducted this analysis in HeLa cells transduced with a whole-genome sgRNA library that utilizes two dual-guide constructs per gene target, with each guide targeting different regions of the same gene (Kirby et al., 2026; Walton et al., 2025) (**Table S5**). At 78 hours following KO induction, we observed a strong correlation with the essential-gene shake-off analysis (**Fig 3A**). This includes the enrichment of gene targets with roles in mitosis, vesicle trafficking, actin organization, and cytokinesis, highlighting the reproducibility of this approach (**Fig 3A**). In addition to essential gene hits, the genome-wide analysis identified other factors such as WASH6P, a regulator of actin polymerization (**Fig 3B**). Although these genes are not fitness-conferring in growth-based assays (DepMap, 2025), this analysis indicates that they contribute to cellular adhesion.

**Figure 3:**
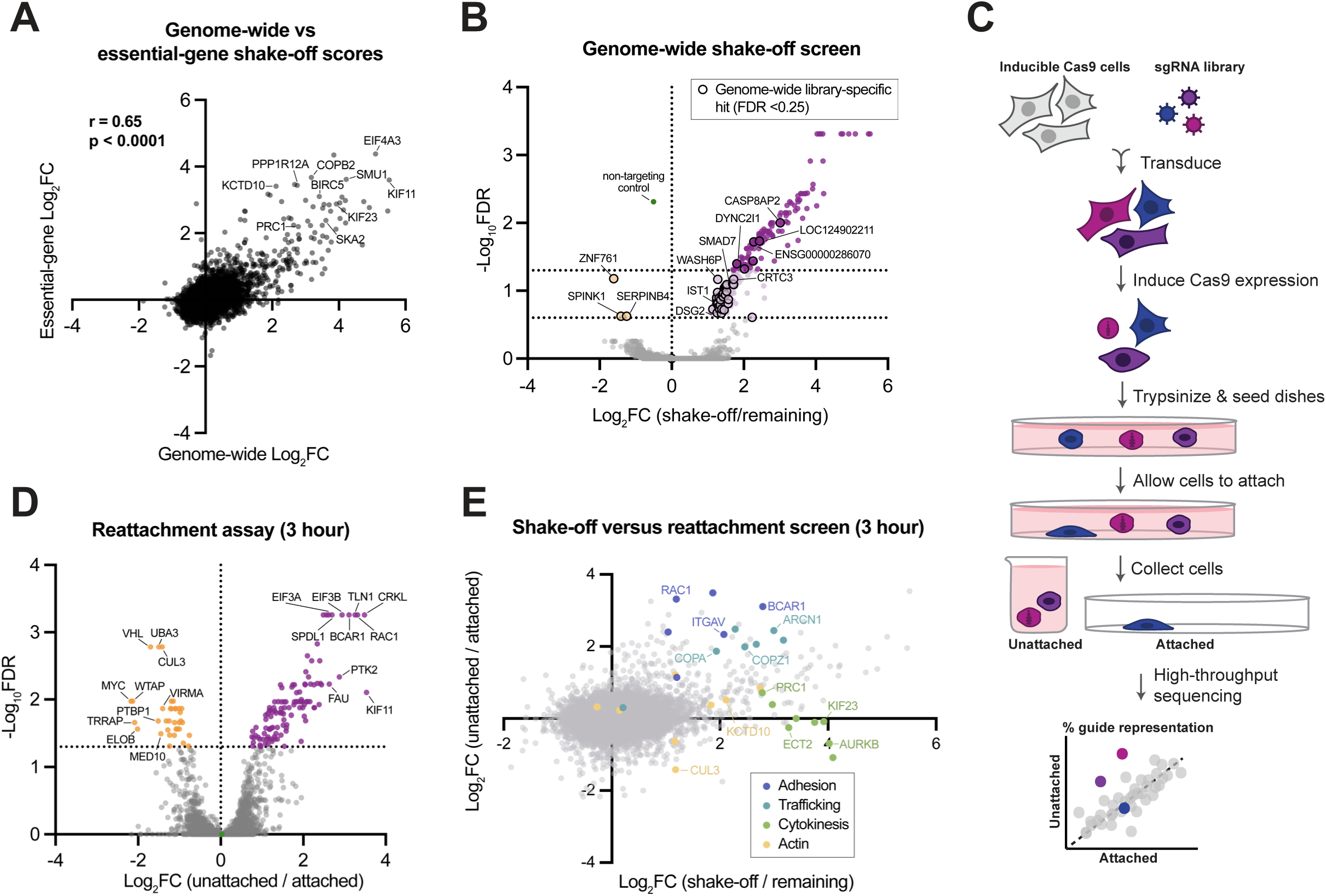
Genome-wide analysis of cellular adhesion via shake-off and reattachment screens. a.) Shake-off scores for genes included in both the genome-wide and essential-gene screens (78-hour and 72-hour knockout induction, respectively). Pearson correlation was performed to determine correlation between scores. b.) Volcano plots of genome-wide shake-off screen in HeLa cells, showing the LFC for each targeted gene in the “shake-off” relative to the remaining cell population following a 78-hour induction. FDR values were derived from MAGeCK enrichment analysis (positive LFC) or depletion analysis (negative LFC). A line is drawn at FDR = 0.05 and FDR = 0.25 to highlight the significantly enriched or depleted genes. Genes targeted in the genome-wide library only (absent from the essential-gene library) with FDR < 0.25 are outlined, and representative genes are labeled. c.) Schematic outlining reattachment screening approach. Cells expressing Cas9 under an inducible promoter were transduced with an sgRNA library. The cells were then seeded onto tissue culture dishes and treated with DOX to induce knockout for ∼78 hours. Cells were trypsinized, reseeded into dishes, and allowed to attach for a set period of time. The unattached cells and the attached cells were each collected and sequenced, and the log_2_ fold change was determined in the unattached population relative to the attached population. d.) Volcano plots of genome-wide reattachment screens in HeLa cells, showing the LFC for each targeted gene in the unattached population relative to the attached cell population following a ∼78-hour induction, collected 3 hours post-reseeding. FDR values were derived from MAGeCK enrichment analysis (positive LFC) or depletion analysis (negative LFC). A line is drawn at FDR = 0.05 to highlight the significantly enriched or depleted genes. Up to 10 significant genes (FDR < 0.05) with the strongest LFC are labeled in each direction. e.) Comparison of the log_2_ fold change between the genome-wide shake-off screen and reattachment screen (using 3-hour reattachment period). Categorized genes from Fig 2C are colored, and select genes are labeled.

For both the functional and morphological screens, cells were given at least 2 days to form stable adhesions prior to the shake-off such that this analysis evaluates the ability of cells to retain adhesion. However, cells also need to first attach to the cell surface before subsequently maturing these more stable adhesions. Thus, to complement the shake-off analysis, we developed a large-scale strategy to assess the initial formation of cellular adhesions (**Fig 3C**). For these experiments, we induced Cas9 expression for 78 hours in HeLa cells transduced with the genome-wide sgRNA library. We then trypsinized cells and re-plated them, allowing cells to re-attach to the plate. Finally, we collected the unattached and attached populations following either 75 minutes or 3 hours post-seeding. At both time points, established adhesion regulators, such as RAC1, TLN1, and PTK2, were among the top hits, validating our approach (**Fig 3D, Fig S3A, Table S5**). Comparison of the results from genome-wide shake-off and attachment screens revealed that vesicle trafficking components were enriched in both analyses, consistent with a requirement for secretory components to deliver adhesion molecules to the cell surface (**Fig 3E, Fig S3B**). In contrast, KCTD10 was identified in the shake-off analysis, but not in the reattachment assay at either time point (**Fig 3E, Fig S3B**). Taken together with the reduced rate of cellular spreading (**Fig 2H**), this suggests that KCTD10 knockout cells are able to attach to the substrate initially but have a reduced ability to mature and maintain adhesion. Finally, as described below, we observed a clear enrichment of established cytokinesis players in the shake-off analysis, but not in the reattachment assay (**Fig 3E, Fig S3B**). Thus, this large-scale approach allowed us to distinguish the requirements to form initial attachments vs. maintain stable substrate adhesion.

### Cells that fail cytokinesis exhibit altered actin organization and behaviors

As described above, gene targets required for the proper completion of cytokinesis strongly perturb cellular adhesion in the shake off assay but not the reattachment assay, suggesting that these genes are required to maintain proper cell-substrate adhesion. Due to the unexpected connection between cytokinesis and cellular adhesion, we next sought to define the basis for this defect. Our shake-off analysis identified each of the established molecular players required for cytokinesis, despite the fact that these gene knockouts do not result in a mitotic arrest (**Fig S2C; Fig 2C**) (Funk et al., 2022). These cytokinesis factors reflect diverse molecular functions, including the regulatory chromosomal passenger complex (AURKB, BIRC5, CDCA8), factors required for spindle midzone organization (PRC1, KIF23), and players cleavage furrow specification and contraction (RACGAP1, ECT2) (Glotzer, 2005). This also includes factors that disrupt cytokinesis indirectly, such as by preventing sister chromatid separation (ESPL1/Separase), which creates a physical block to cytokinesis by trapping DNA in the cleavage furrow (Hauf et al., 2001). This range of diverse factors suggests that the inability to complete cytokinesis causes the adhesion defect instead of reflecting a secondary role for some proteins in adhesion. To evaluate the role of the successful completion of cytokinesis in modulating cell-substrate adhesion, we first examined cytoskeletal morphology in knockouts of ECT2, a RhoGEF required for contractile ring formation in cytokinesis, and PRC1, a microtubule-binding component of the central spindle important for defining the division plane, on fibronectin-coated surfaces. To mark the cell body and ensure the proper identification of multinucleated cells, we transfected the cells with the membrane mem-mCh marker (Kiyomitsu and Cheeseman, 2012). In cells that had failed cytokinesis, microtubule organization appeared largely normal (**Fig 4A**). In contrast, we observed a dramatic change in the organization of the actin cytoskeleton. We often observed cases in which the portion of the cell containing the nuclei was devoid of actin, whereas the other portion of the cell displayed bright actin stress fibers (**Fig 4A, Fig S4A**). Indeed, quantification of actin distribution revealed a reduction of actin surrounding the nucleus and at the cortex (**Fig 4A**).

**Figure 4:**
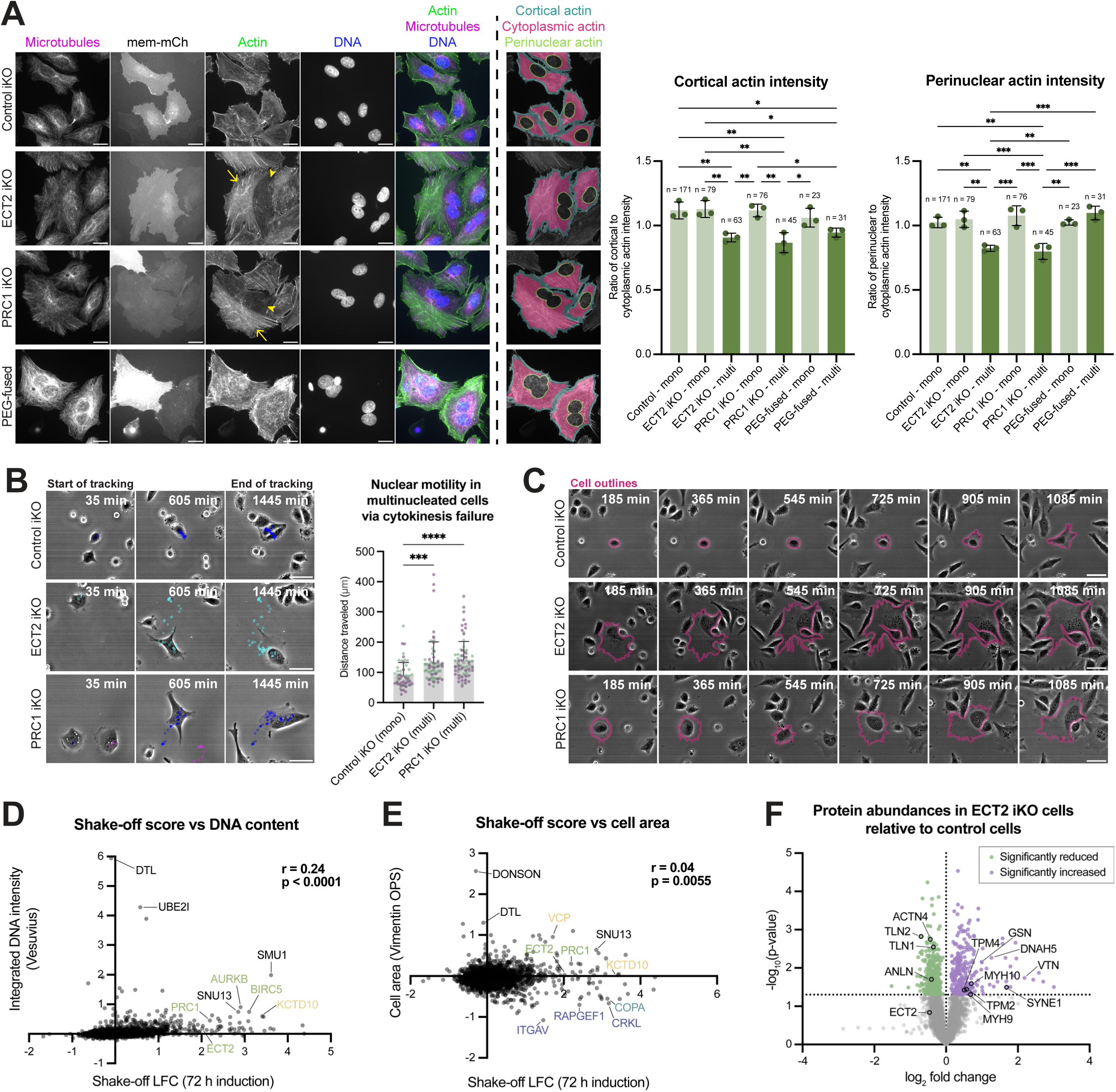
Cells that fail cytokinesis display altered actin organization and nuclear motility. a.) (Left) Representative z-projected immunofluorescence images of control, ECT2 iKO, PRC1 iKO, and PEG-fused control iKO cells following a 72-hour induction, seeded on fibronectin-coated coverslips. Actin (phalloidin), microtubules (anti-α Tubulin), mem-mCherry, and DNA (Hoechst). Arrows and arrowheads point to stress fiber-containing and actin-devoid regions, respectively. Scale bars, 20 µm. Representative CellProfiler overlays of the cortical, cytoplasmic, and perinuclear regions used for quantification are shown. (Right) Quantification of perinuclear or cortical actin intensity relative to cytoplasmic intensity. Total number of cells quantified across all reps are noted. Data represent mean ± SD. One-way ANOVA was performed (\**p* ≤ 0.05, \*\**p* ≤ 0.01, \*\*\**p* ≤ 0.001; \*\*\*\**p* ≤ 0.0001, nonsignificant p-values are not shown). b.) (Left) Representative brightfield images with traces of nuclear positions in control, ECT2, and PRC1 iKO HeLa cells acquired in the spreading assay, where cells were trypsinized and re-plated onto a fibronectin-coated surface and imaged over time. Time following the start of imaging is indicated. Scale bars, 50 µm. (Right) Quantification of nuclear motility, measured as the total distance travelled by nuclei. For conditions that produce multinucleated cells, only multinucleated cells were tracked. Color indicates experimental replicate. Data represent mean ± SD. One-way ANOVA was performed (\**p* ≤ 0.05, \*\**p* ≤ 0.01, \*\*\**p* ≤ 0.001; \*\*\*\**p* ≤ 0.0001, nonsignificant p-values are not shown). c.) Representative brightfield images of control, ECT2, and PRC1 iKO HeLa cells acquired in the spreading assay, where cells were trypsinized and re-plated onto a fibronectin-coated surface and imaged over time. Time following the start of imaging is indicated. Cells are manually outlined for visualization. Scale bars, 50 µm. d.) Shake-off scores (log_2_ fold change) from the essential-gene screen in HeLa (72-hour induction) versus nuclear integrated DNA intensity from Vesuvius (78-hour induction). Select points are labeled and colored corresponding to phenotypic clusters in Fig 2C. Pearson correlation was performed to determine correlation between scores. e.) Shake-off scores (log_2_ fold change) from the essential-gene screen in HeLa (72-hour induction) versus cell area from Vimentin OPS (78-hour induction). Select points are labeled and colored corresponding to phenotypic clusters in Fig 2C. Pearson correlation was performed to determine correlation between scores. f.) Volcano plot comparing protein abundance in control and ECT2 iKO HeLa cells as measured by quantitative TMT-MS. Proteins with significant changes in abundance as determined by a Student’s t-test (p < 0.05) are highlighted. Select proteins are labeled.

Given the substantial change in actin organization in cells that have failed cytokinesis, we next considered the adhesion dynamics of these cells. In the cell spreading assay, we did not observe reduced spreading rates for either the ECT2 or PRC1 knockout (**Fig S4B, Fig S4C**). However, we observed a striking difference in the dynamics of ECT2 and PRC1 knockout cells. In particular, although HeLa cells are not motile, multinucleated cells that had failed cytokinesis displayed markedly enhanced movement when marked by nuclear position (**Fig 4B)**. This increased motility appears to be caused in part by a unique phenotype in which nuclei cluster together and migrate around the cell, occasionally driving cell movement (**Fig 4B**, **Fig 4C**). This suggests that failed cytokinesis does not disrupt the initial attachment of cells and adhesion formation, but that these cells display altered actin and adhesion dynamics that may be responsible for their enrichment in the shake-off analysis.

We next considered whether these altered behaviors and reduced adhesion are a consequence of having a larger cell size or increased DNA content. Based on the analysis of quantitative morphological parameters derived from the Vesuvius and Vimentin optical pooled screening datasets, shake-off score is only weakly associated with integrated DNA intensity and is not correlated with cell size (**Fig 4D**, **Fig 4E**). Indeed, knockouts that resulted in the largest increase in cell size or DNA content (e.g., DTL) did not exhibit increased susceptibility to shake-off. To evaluate whether cytokinesis failure itself or the resulting polyploidy influences cellular adhesion, we generated artificially multinucleated cells using PEG-mediated cell fusion and monitored cell spreading dynamics. Similarly to cells that have failed cytokinesis, PEG-fused cells do not display a noticeable reduction in the rate of cell spreading (**Fig S4D**) and also exhibit an increase in nuclear motility (**Fig S4E**). However, we rarely observed the same nuclear clustering phenotype (**Fig S4F**). In addition, although the actin cytoskeleton appears altered in PEG-fused cells, these cells did not exhibit a similar loss of perinuclear or cortical actin or the absence of actin in one portion of the cell (**Fig 4A**, **Fig S4A**). Taken together, this suggests that the altered adhesion and motility observed in cytokinesis knockouts are due in part to multinucleation, while some aspects reflect cytokinesis-failure specifically.

Finally, to understand the molecular mechanisms underlying these phenotypes, we performed quantitative proteomics on ECT2 iKO cells (**Fig 4F**, **Table S6**). Amongst the most dramatically upregulated proteins was the LINC complex component, Nesprin-1 (SYNE1), which is more than 3-fold more abundant in ECT2 iKO cells compared to controls. The LINC complex connects the nucleus to the actin cytoskeleton and is required for nuclear positioning (Gundersen and Worman, 2013). We also found increased levels of Lamin-A/C (LMNA), which would be predicted to increase nuclear stiffness (Swift et al., 2013), as well as increased levels of the myosin subunits MYH9 and MYH10 and tropomyosins TPM2 and TPM4. As we found that reduced MYH9 resulted in increased ability to resist detachment with mechanical shake-off (**Fig 1B**), increases in MYH9 are predicted to increase contractility and reduce adhesion. Finally, we identified a reduction of focal adhesion-associated proteins, including talins (TLN1 and TLN2) and α-actinin-4 (ACTN4). The combination of these changes suggests that ECT2 iKO cells have stiffer nuclei with increased cytoskeletal coupling and contractility, which could be responsible for the altered nuclear dynamics, lending the cell to a mechanically-unstable and detachment-prone state.

Taken together, this work implicates a requirement for successful cytokinesis to maintain cytoskeletal integrity and resist detachment with mechanical pressure. In addition to providing novel insights into cellular adhesion, this work has substantial implications for cancer metastasis, where tetraploidy has been linked to cancer cell migration and invasiveness (Fujiwara et al., 2005; Ganem et al., 2007; Jemaa et al., 2017; Jemaa et al., 2023).

## Acknowledgements

This work was supported by grants from the NIH (R35GM126930 to I.M.C and P01AI120943-06A1, RM1NS133601, and R61CA278536 to P.B.) and Chan Zuckerberg Initiative grants (Cell Biology @ Scale) to I.M.C. and P.B. M.D. is supported in part by an NSF GRFP fellowship (000955563). This research was also supported by the Whitehead Innovation Initiative (to I.M.C.). We thank the Whitehead Quantitative Proteomics Core for mass spectrometry, the Whitehead Flow Cytometry Core for cell sorting, the Whitehead Genome Technology Core for library sequencing, the Whitehead Functional Genomics Platform for assistance with screen design execution, and the Whitehead Bioinformatics & Research Computing for screen analyses. We thank the members of the Cheeseman lab for helpful discussions. We thank the Broad Institute’s Genetic Perturbation Platform for supporting the construction of the genome-wide dual-guide library.

## Supplemental Figure Legends

**Supplemental Figure 1:**
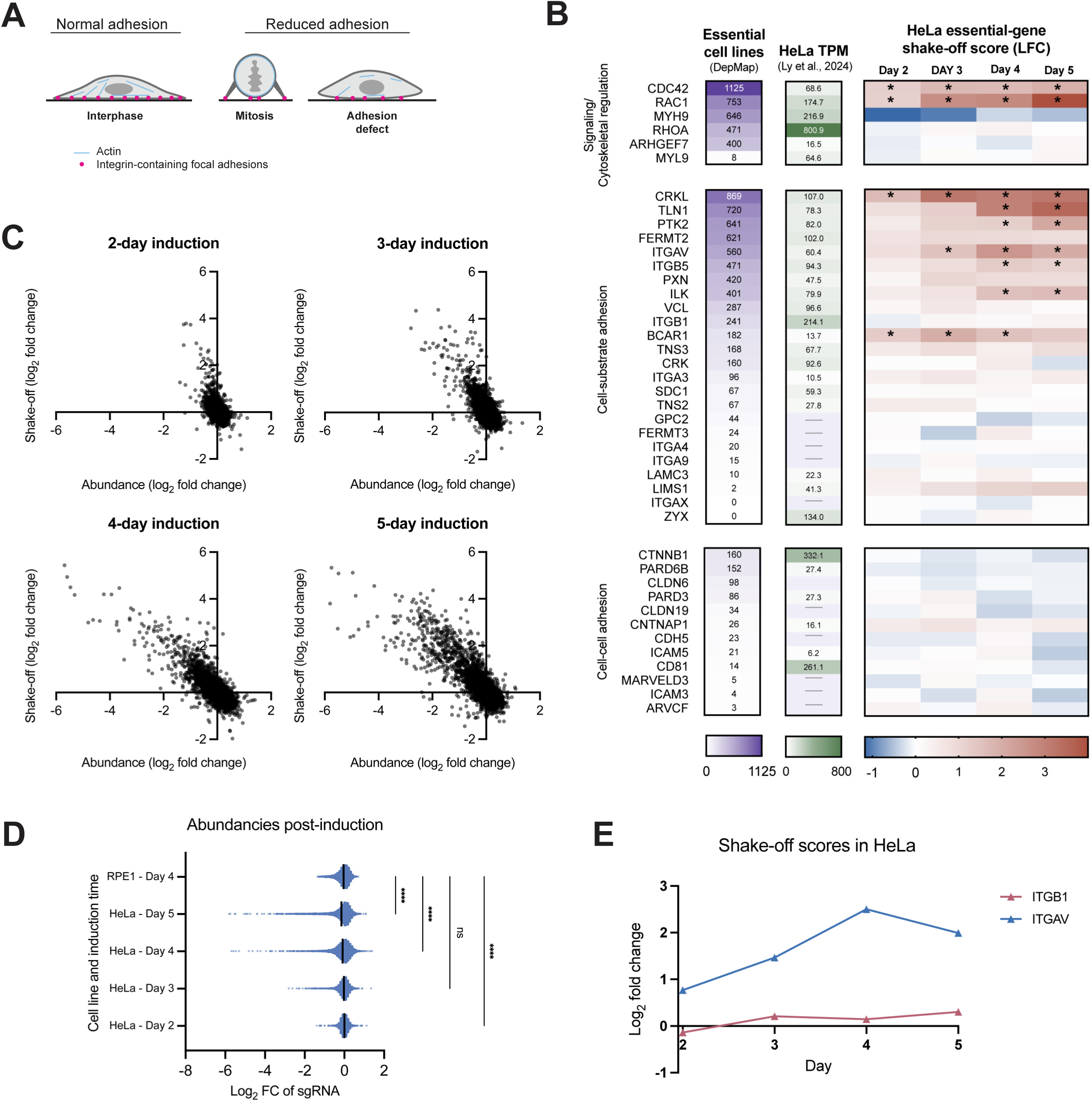
Supporting analyses of essential-gene shake-off screens. a.) Cartoon highlighting factors contributing to adhesion strength in cultured cells. b.) Known adhesion genes present in essential gene library and the number of cell lines in which each gene is deemed essential (DepMap), expression level in HeLa cells (Transcripts Per Million (TPM)), and shake-off scores (log_2_ fold change) from the essential-gene screens in HeLa. Asterisks indicate FDR < 0.05 from MAGeCK enrichment analysis. c.) Gene-level sgRNA depletion score (LFC, remaining vs. day 0) versus shake-off score (LFC, shake-off vs. remaining) following 2, 3, 4, or 5 days of Cas9 induction in HeLa cells transduced with an essential-gene sgRNA library. d.) Bar graph of log_2_ fold change in gene-level sgRNA abundance at the indicated induction time relative to the initial population. Line indicates the mean. One-way ANOVA was performed (ns, not significant; \**p* ≤ 0.05, \*\**p* ≤ 0.01, \*\*\**p* ≤ 0.001; \*\*\*\**p* ≤ 0.0001). e.) Shake-off scores for ITGB1 and ITGAV in HeLa essential-gene screen at different time points following Cas9 induction.

**Supplemental Figure 2:**
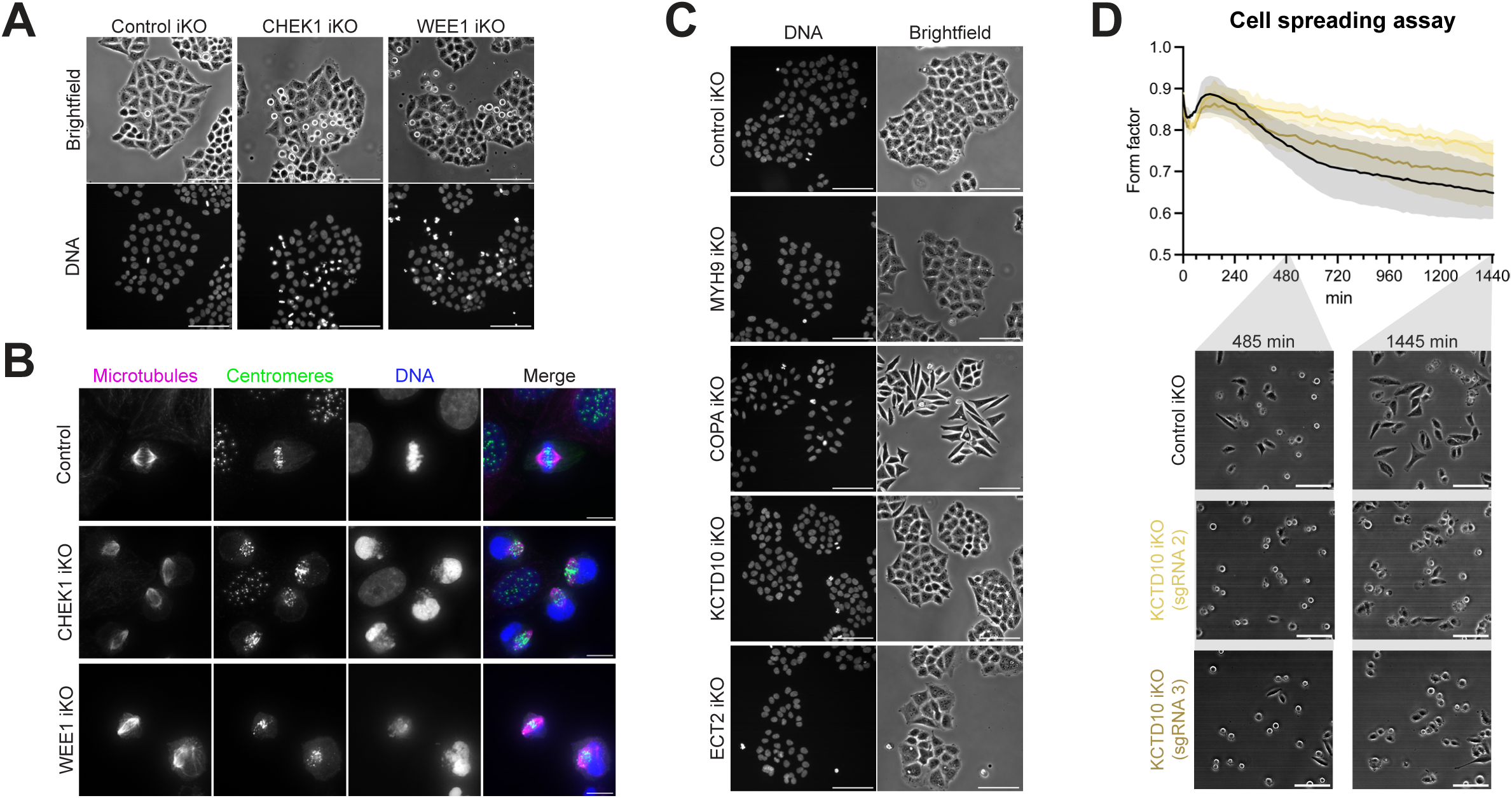
Further validation of hits from shake-off screens. a.) Representative brightfield and fluorescent images of live control, CHEK1, and WEE1 iKO HeLa cells following 72-hour induction. DNA (Hoechst). Scale bar, 100 µm. b.) Representative z-projected deconvolved immunofluorescence images of control, CHEK1, and WEE1 iKO HeLa cells following 72-hour induction. Microtubules (anti-alpha tubulin), centromeres (ACA), and DNA (Hoechst). Scale bars, 10 µm. c.) Representative brightfield and fluorescent images of live control MYH9, COPA, KCTD10, and ECT2 inducible knockout cells following 72-hour induction. DNA (Hoechst). Scale bar, 100 µm. d.) Cell spreading assay in control and KCTD10 iKO HeLa cells expressing different sgRNAs. Spreading assays with iKO cells were performed in parallel; the same control data are shown in each figure panel for comparison. (Top) Quantification of cell form factor of cells over time following trypsinization and re-plating onto fibronectin-coated surfaces, averaged across two experimental replicates (*n* > 150 per condition per replicate). Data represent mean ± SEM. (Bottom) Representative brightfield images of cells at select time points. Scale bars, 100 µm.

**Supplemental Figure 3:**
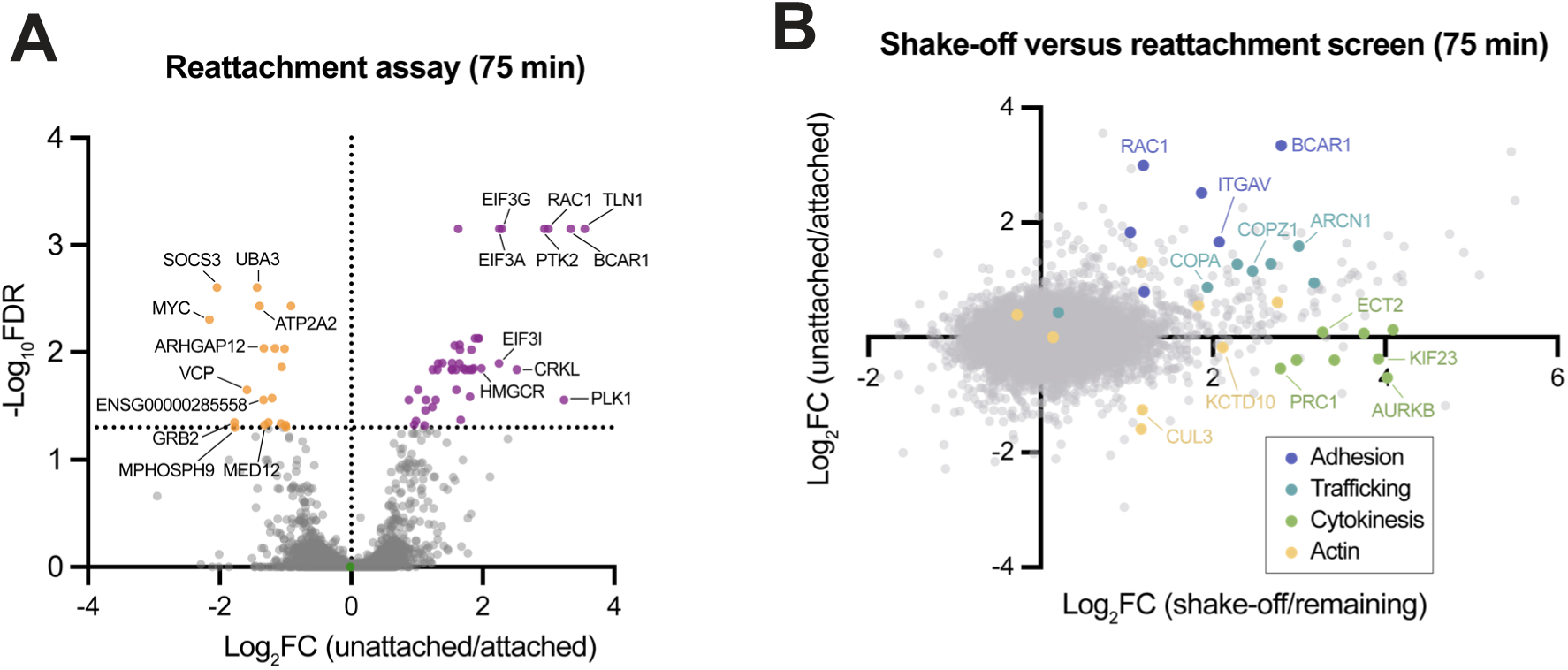
Reattachment screen following a 75-minute reattachment period. a.) Volcano plots of genome-wide reattachment screens in HeLa cells, showing the LFC for each targeted gene in the unattached population relative to the attached cell population following a ∼78-hour induction, collected 75 minutes post-reseeding. FDR values were derived from MAGeCK enrichment analysis (positive LFC) or depletion analysis (negative LFC). A line is drawn at FDR = 0.05 to highlight the significantly enriched or depleted genes. Up to 10 significant genes (FDR < 0.05) with the strongest LFC are labeled in each direction. b.) Comparison of the log_2_ fold change between the genome-wide shake-off screen and reattachment screen (using 75-minute reattachment period). Categorized genes from Fig 2C are colored, and select genes are labeled.

**Supplemental Figure 4:**
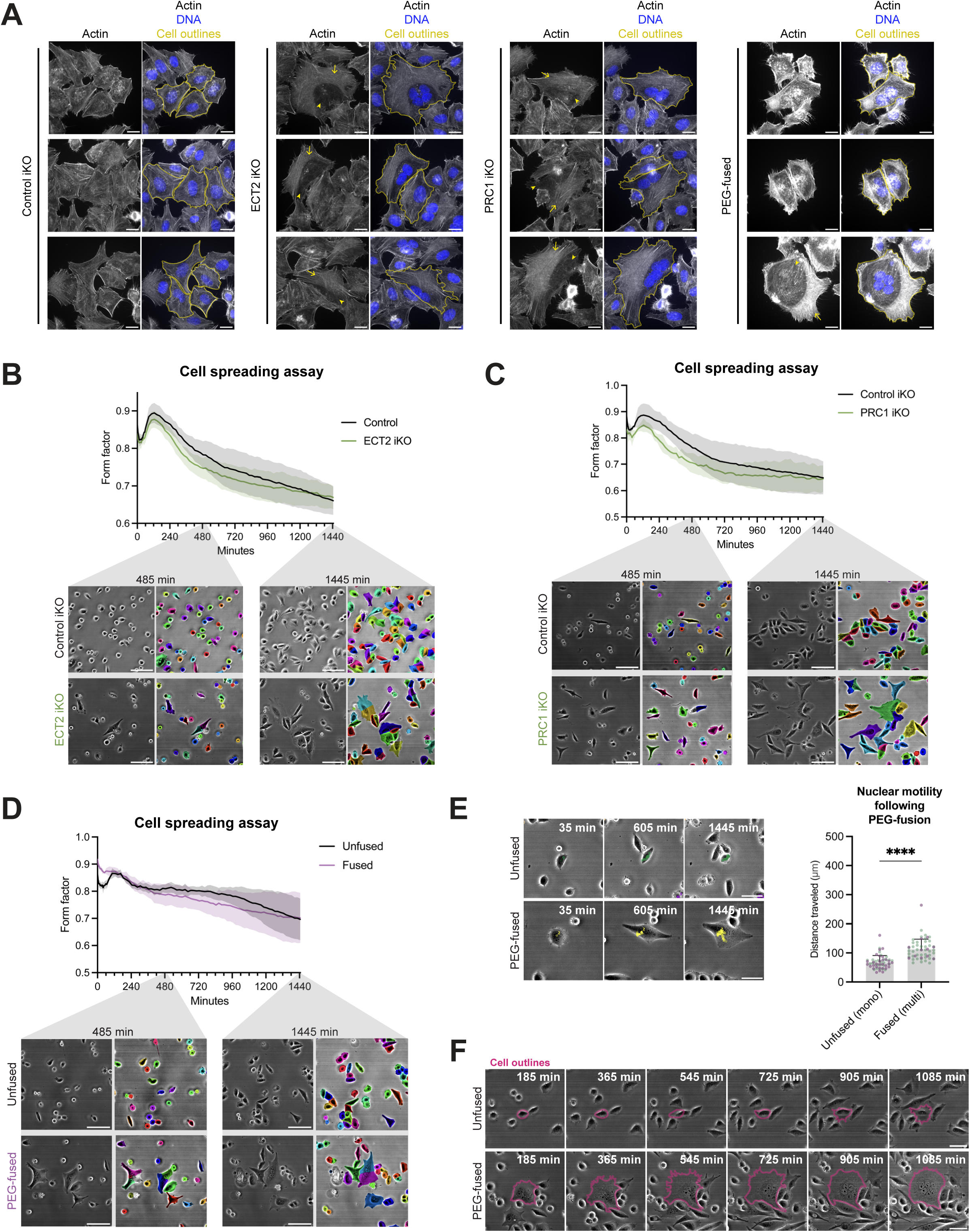
Supporting results for the analysis of the role of successful cytokinesis in adhesion. a.) Additional representative z-projected immunofluorescence images of control, ECT2 iKO, PRC1 iKO, and PEG-fused control iKO cells following a 72-hour induction, seeded on fibronectin-coated coverslips. Actin (phalloidin) and DNA (Hoechst). Arrows and arrowheads point to stress fiber-containing and actin-devoid regions, respectively. Cells are manually outlined for visualization. Scale bars, 20 µm. b.) Cell spreading assay in control and ECT2 iKO HeLa cells. Spreading assays with iKO cells were performed in parallel; the same control data are shown in each figure panel for comparison. (Top) Quantification of cell form factor of cells over time following trypsinization and re-plating onto fibronectin-coated surfaces, averaged across three experimental replicates (*n* > 150 per condition per replicate). Data represent mean ± SEM. (Bottom) Representative brightfield images and Cellpose-SAM segmentation of cells at select time points. Scale bars, 100 µm. c.) Cell spreading assay in control and PRC1 iKO HeLa cells. Spreading assays with iKO cells were performed in parallel; the same control data are shown in each figure panel for comparison. (Top) Quantification of cell form factor of cells over time following trypsinization and re-plating onto fibronectin-coated surfaces, averaged across two experimental replicates (*n* > 150 per condition per replicate). Data represent mean ± SEM. (Bottom) Representative brightfield images and Cellpose-SAM segmentation of cells at select time points. Scale bars, 100 µm. d.) Cell spreading assay in unfused and PEG-fused HeLa cells. (Top) Quantification of cell form factor of cells over time following trypsinization and re-plating onto fibronectin-coated surfaces, averaged across two experimental replicates (*n* > 150 per condition per replicate). Data represent mean ± SEM. (Bottom) Representative brightfield images and Cellpose-SAM segmentation of cells at select time points. Scale bars, 100 µm. e.) (Left) Representative brightfield images with traces of nuclear positions in unfused and PEG-fused HeLa cells acquired in the spreading assay, where cells were trypsinized and re-plated onto fibronectin-coated surface and imaged over time. Time following the start of imaging is indicated. Scale bars, 50 µm. (Right) Quantification of nuclear motility, measured as the total distance travelled by nuclei. For conditions that produce multinucleated cells, only multinucleated cells were tracked. Color indicates experimental replicate. Data represent mean ± SD. Student’s t-test was performed (\**p* ≤ 0.05, \*\**p* ≤ 0.01, \*\*\**p* ≤ 0.001; \*\*\*\**p* ≤ 0.0001, nonsignificant p-values are not shown). f.) Representative brightfield images of unfused and PEG-fused HeLa cells acquired in the spreading assay, where cells were trypsinized and re-plated onto fibronectin-coated surface and imaged over time. Time following the start of imaging is indicated. Cells are manually outlined for visualization. Scale bars, 50 µm.

## Supplemental tables

**Table S1: Essential-gene shake-off screen sgRNA sequences, counts, and gene scores.**

**Table S2: Gene set enrichment analysis (GSEA) of essential-gene shake-off screens in HeLa and Rpe1 cells (enrichment in “shake-off”).**

**Table S3: Results from optical pooled screening with α-Vimentin, including feature table and clustering.**

**Table S4. TMT MS analysis of KCTD10 vs control iKO.**

**Table S5: Genome-wide shake-off and reattachment screen sgRNA sequences, counts, and gene scores.**

**Table S6: TMT MS analysis of ECT2 vs control iKO. Table S7: List of all cell lines used in this study.**

## Methods

## Key Resource Table

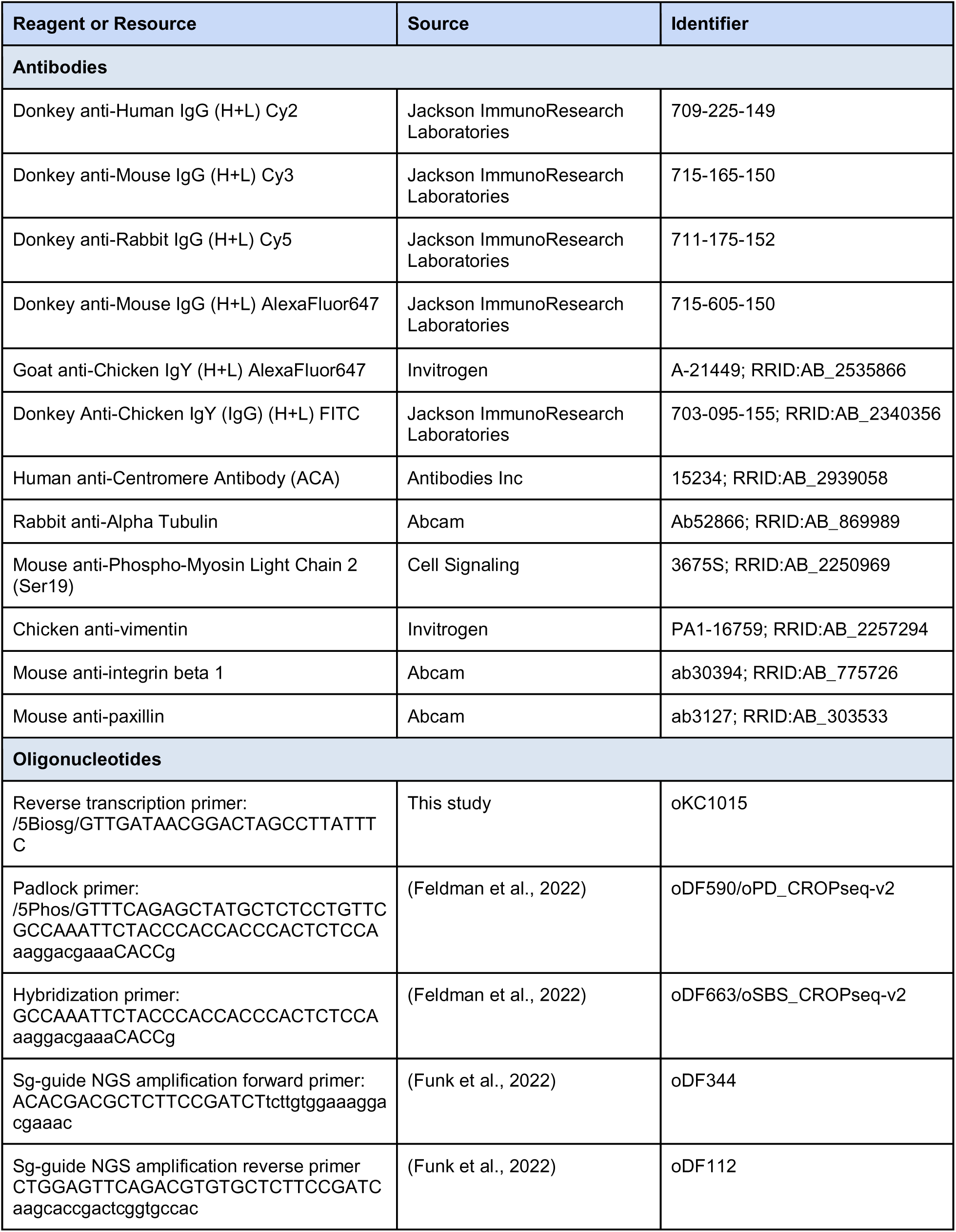
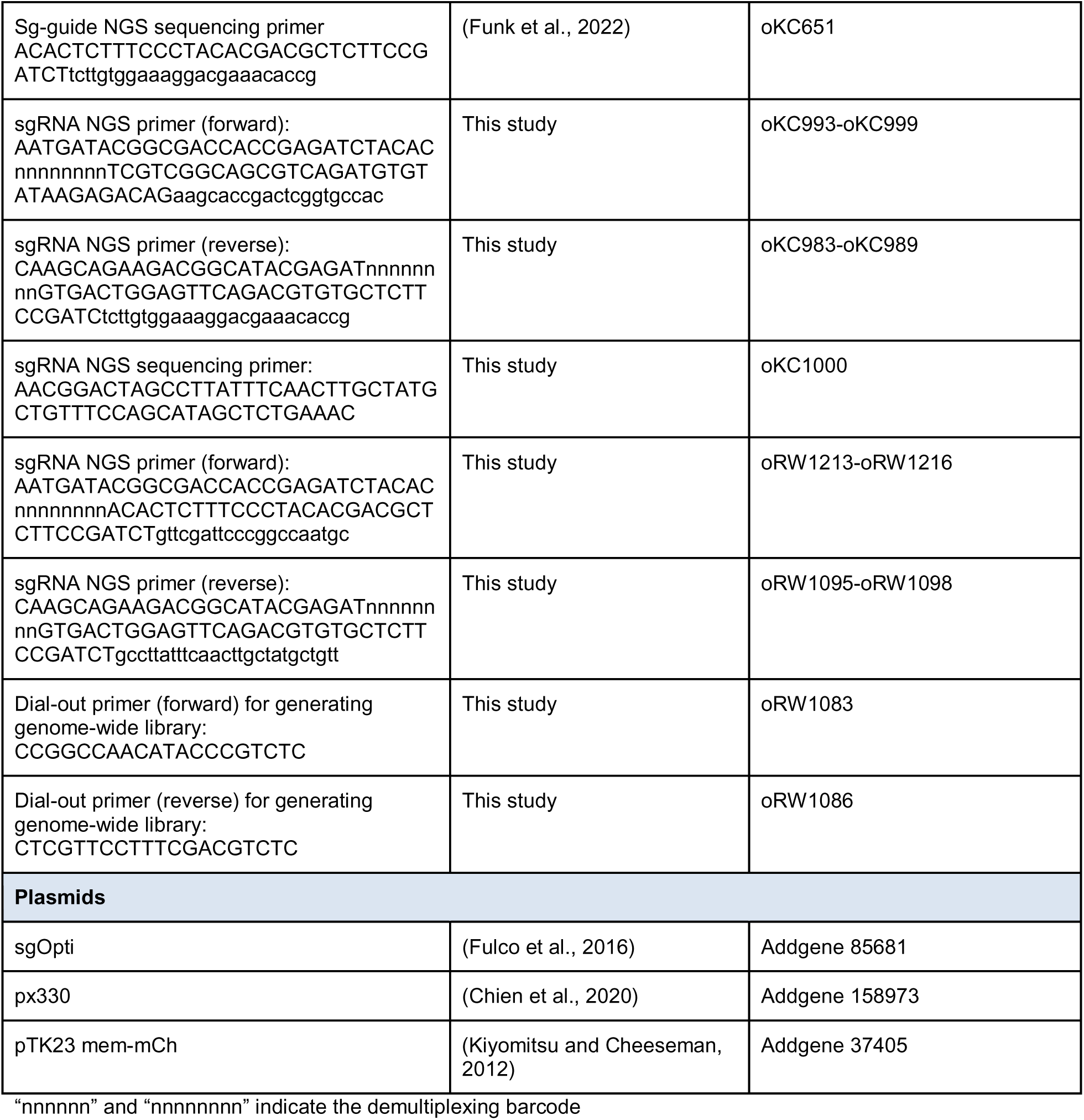

### Tissue culture

HeLa and RPE1 cells were cultured in Dulbecco’s modified Eagle medium (DMEM) supplemented with 10% fetal bovine serum (FBS), 100 U/mL penicillin and streptomycin, and 2 mM L-glutamine at 37°C with 5% CO_2_. Doxycycline-inducible cell lines were cultured in medium containing tetracycline-free FBS, and were induced with 1 µg/mL doxycycline hyclate. Doxycycline-containing media was renewed every ∼24 hours. Cells were routinely monitored for mycoplasma contamination using commercial detection kits.

### Generation of cell lines

Cell lines used in this study are listed in Table S7. To generate single inducible knockout cell lines, sgRNAs were cloned into the sgOpti plasmid (puromycin-resistant, Addgene #85681) and introduced into an inducible Cas9 HeLa cell line (McKinley et al., 2015) via lentiviral transduction (Wang et al., 2015). Transduced cells were selected with 0.5 μg/ml puromycin to generate polyclonal cell lines.

To generate a puromycin-sensitive, inducible Cas9 RPE1 cell line (cKC580), cTT33.1 (McKinley and Cheeseman, 2017) were transiently transfected with Cas9 and sgRNAs removing puromycin resistance (GCTCGGTGACCCGCTCGATG and GTGGTCGGGCGAGACGCCGA) using px330 (Chien et al., 2020).

For expression of mem-mCh, pTK23 (Addgene #37405) (Kiyomitsu and Cheeseman, 2012) was transiently transfected into cells using Xtremegene-9 (Roche) ∼48 hours prior to analysis.

### Generation of cell libraries for pooled CRISPR screens

Two separate sgRNA libraries were used for pooled CRISPR screens in this study. The essential-gene sgRNA plasmid library was generated in a previous study, and is composed of 20,445 sgRNAs in the CROPseq-puro-v2 vector (Feldman et al., 2022; Funk et al., 2022). The dual-cutting genome-wide library was generated as a part of this study, described as follows.

We designed a dual-guide CROPseq-multi library to target all 20,553 genes (the union of protein-coding genes from GENCODE, RefSeq, and CHESS). We designed 4 unique guides per gene using CRISPick (Doench et al., 2016; Drepanos et al., 2026; Sanson et al., 2018) and paired the guides according to computational rankings (“Pick Order” 1+4 for construct 1 and 2+3 for construct 2) for a total of two constructs per gene. For targets with only three (129 genes) or two (117 genes) guides passing the design criteria, the top two guides were paired and represented twice in the library with unique internal barcodes (iBARs). For targets with only one guide (95 genes) passing the design criteria, the gene-targeting guide was paired with a nontargeting guide and represented twice in the library with unique iBARs. We additionally included 600 intergenic cutting controls and 200 nontargeting controls. Intergenic cutting controls tile all chromosomes and match the distribution of gene-targeting guide pairs with respect to on- and off-target predictions and inter-guide-pair distance. Guide pairs were randomly assigned one of three orthogonal tRNAs in equal proportions. Each guide was assigned a 12 nucleotide iBAR sequence. The iBAR library has Levenshtein distance 3 between all barcodes to enable error correction. All combinations of guide position (1 or 2) and iBAR are unique. Some iBARs are used in both guide positions with the guide position distinguished by the orthogonal sgRNA scaffold sequences. Two sub-libraries were constructed: one containing all gene-targeting guides and controls (41,906 dual-guide constructs) and a second dedicated negative control library encoding nontargeting controls, intergenic controls, and 412 olfactory receptor genes (1,624 dual-guide constructs). The separate control library encodes identical constructs represented in the complete library and is used as a “spike-in” library to flexibly increase control representation.

Oligo pools (Twist Biosciences) were cloned into plasmid libraries as previously described (Walton et al., 2025). Briefly, oligo pools were amplified with 1X KAPA HiFi HotStart Ready Mix (Roche KK2601), 1X EvaGreen qPCR dye (Biotium 31000), 1M Betaine (MilliporeSigma B0300), 300 nM forward and reverse primers (Integrated DNA Technologies), and 12 pg/µL of template oligo pool in 50 µL reactions. PCRs were conducted with the following thermal cycling protocol: 95 °C for 3 min, 16 cycles of (98 °C for 20 s, 62 °C for 15 s, 72 °C for 15 s), then 72 °C for 1 min. We performed 200 reactions for the full 41,906-member library and 10 reactions for the 1,624-member library. Plasmid libraries were then cloned into CROPseq-multi-v2-Puro (Addgene #225752) by restriction digestion and ligation. Each 20 µL ligation reaction consisted of 10 femtomole (fmol) of BsmBI-digested amplified oligo pool, 20 fmol of BsmBI-digested, dephosphorylated, and gel-purified plasmid backbone, 1X T4 DNA ligase buffer (New England Biolabs M0202S), and 20 U/µL of T4 DNA ligase (New England Biolabs M0202S). We performed 21 ligation reactions for the 41,906-member library and 1 for the 1,624-member library. After overnight ligation at 16°C, assemblies were heat-inactivated, purified with Ampure XP paramagnetic beads (Beckman Coulter A63880), and eluted in water (40 µL for the 41,906-member library and 5 µL for the 1,624-member library). Purified ligations were electrotransformed by combining 5 µL of ligation and 25 µL of Endura electrocompetent cells (Biosearch Technologies 60242-2). We performed 8 transformations for the 41,906-member library and one transformation for the 1,624-member library. We prepared next generation sequencing libraries with a single-step PCR, sequenced on an AVITI 2×150 Sequencing Kit Cloudbreak FS Low Output (Element Biosciences 860-00011), and analyzed using our GitHub package (https://github.com/rtwalton/CROPseq-multi). We validated mapping rates of 78-85% per barcode, uniformity (ratio of the 90th to 10th percentile abundance) of 2.8-2.9, and recombination frequencies of 0.4-0.6%.

### Lentiviral library production and transduction

Lentiviral libraries were prepared by transfecting HEK-293T cells with a mixture containing 20 µg plasmid library, 3.62 µg pCMV-VSV-G, and 8.28 µg psPAX2, 95.8 µL FuGENE 4K (essential-gene) or 76.80 µL Xtremegene-9 transfection reagent (genome-wide), and Opti-MEM media to a final volume of 2 mL (Fugene 4K mix) or 1 mL (Xtremegene-9 mix). Plasmids were mixed at a 19:1 ratio for the 20K essential-gene:250 control plasmid library. Cells were seeded at 15 x 10^6^ cells in a T175 cm^2^ flask in 20 mL DMEM supplemented with 10% fetal bovine serum and 1% penicillin-streptomycin 30 h pre-transfection, and media was changed to 20 mL virus production media (IMDM supplemented with 20% heat-inactivated fetal bovine serum) 7-8 h pre-transfection. Media was changed to 55 mL fresh virus production medium 16-17 h post-transfection, viral supernatant was harvested 48 h post-transfection, 25 mL fresh virus production medium was added, and viral supernatant was harvested at 72 h post-transfection. Viral supernatant was 0.45 µm filtered and stored at -80 °C immediately after harvest.

The essential-gene sgRNA library was introduced into HeLa cells in a prior study (Funk et al., 2022). To introduce the essential-gene library into RPE1 cells, RPE1-TetR-Cas9 (cKC580) cells were transduced with the library via lentiviral transduction and selected with 2 μg/mL puromycin (Millipore; P4512) for 5 days. To introduce the genome-wide library into HeLa cells, HeLa A1 cells were transduced with the genome-wide library virus and selected 2 days later with 2 μg/mL puromycin (Millipore; P4512) for 4 days.

### Shake-off screens

The essential-gene iKO HeLa, essential-gene iKO RPE1, or genome-wide iKO HeLa cell libraries were induced with 1 μg/mL doxycycline hyclate for the indicated amount of time and plated at least 2 days before harvest. Prior to induction, cells were collected and pelleted for DNA extraction and sequencing. Following induction, to enrich for weakly adhered cells, cells in 15 cm^2^ plates were tapped against each other and washed repeatedly with media to dislodge cells. This “shake-off” population along with the cells that remained on the plate (“remaining” population) were collected. Cells were then pelleted for DNA extraction and sequencing. The relative sgRNA abundance was compared between the shake-off and remaining cell populations. The remaining cell population at each induction time point was also compared to the initial cell population to determine gene dropout.

### Reattachment screen

The genome-wide iKO HeLa cell library was cultured and induced with 1 μg/mL doxycycline hyclate for approximately 78 hours. Prior to induction, cells were collected and pelleted for DNA extraction and sequencing. Following induction, cells were detached from plates by minimal incubation with 0.05% trypsin/EDTA at 37°C. Trypsinized cells were spun down, resuspended in pre-warmed media, and plated in 15 cm^2^ plates at a density of 15 million cells per plate. After 75 minutes or 3 hours, attached and unattached cell populations were collected. To collect the “unattached” population, the media was collected. Plates were gently washed with PBS two times and the PBS washes were pooled with the media. To collect the “attached” population, cells that remained on the plate were detached with trypsin. Both populations were then pelleted for DNA extraction and sequencing.

### Sequencing library preparation for shake-off and reattachment screens

For screens using the essential-gene library in HeLa cells, genomic DNA was extracted using PureLink (Invitrogen). sgRNA sequences were then PCR amplified using Q5 hotstart (NEB) with primers oDF344 and oDF112 before addition of index barcodes and sequencing using an Illumina sequencer using sequencing primer oKC651. For remaining screens, genomic DNA was extracted using Blood genomicPrep Mini Spin (Cytiva) or QIAmp DNA Blood Maxiprep (Qiagen) kit. ExTaq DNA Polymerase (Takara Bio) was used to amplify the guide sequence using indexed primers (oKC993-999 and oKC983-oKC989 for the essential-gene library in RPE1; oRW1213-oRW1216 and oRW1095-oRW1098 for the genome-wide library in HeLa). Custom primer oKC1000 was used to sequence the essential-gene RPE1 library.

### Screen analysis for shake-off and reattachment screens

For screens using the essential-gene library, sequences were trimmed and mapped to the sgRNA library using Bowtie v1.0.0 or v1.2.2.

For screens using the dual-guide genome-wide library, paired-end 150 bp reads were assigned to library constructs following the library_NGS_analysis.ipynb notebook from CROPseq-multi (https://github.com/rtwalton/CROPseq-multi), with max_reads=1e9 to accommodate our large genome-wide sequencing datasets. We modified the pipeline to permit up to 1 mismatch per barcode element (max_mismatches=1). Specifically, for each barcode element (spacer_1, iBAR_1, spacer_2, iBAR_2), a 1-mismatch look-up table was constructed containing all single-nucleotide variants of each reference sequence, excluding variants shared by multiple references. For each element, an exact match to a reference sequence was accepted directly; only in the absence of an exact match was the look-up table consulted. A barcode element found in the look-up table was corrected to its corresponding reference sequence. A read pair was assigned a count to a library construct only if all four barcode elements were successfully matched — either exactly or via mismatch correction — and arranged in the expected order (spacer_1–iBAR_1–spacer_2–iBAR_2).

The MAGeCK RRA v0.5.9.5 algorithm (Li et al., 2014) test was used to analyze each comparison separately using the following parameters: Adjustment method: FDR; FDR threshold: 0.05; Log2FC method: mean.

### GSEA analysis

Gene scores representing the mean log2 fold change in abundance of the sgRNAs targeting a given gene were plotted and statistical analyses were performed using R or GraphPad Prism. Pre-ranked gene set enrichment analysis of CRISPR-Cas9 screens were performed in the GSEA program using mean log2 fold change gene scores with the classic normalization scheme, meandiv normalization method, absolute max of probes mode, minimum and maximum gene set sizes of 15 and 500, respectively, and 1000 permutations with the Human_Gene_Symbol_with_Remapping_MSigDB.v2025.1.Hs.chip gene name file and the following gene set file: c5.go.cc.v2025.1.Hs.symbols.gmt.

### Optical pooled screen

Optical pooled screens were carried out as previously described (Feldman et al., 2022; Funk et al., 2022). The essential-gene inducible KO HeLa cell library was induced with 1 μg/mL doxycycline hyclate for 78 hours and seeded at a density of 350k cells/well in 6-well glass-bottom plates (Cellvis, P06-1.5H-N) 48 hours prior to fixation. Cells were fixed with 4% formaldehyde (Electron Microscopy Sciences, 15710) and 0.007% glutaraldehyde (v/v) in PBS for 25 minutes. All steps used RNase free reagents to preserve mRNA integrity up until the reverse transcription step was complete. The cells were then washed with PBS and permeabilized with 70% ethanol for 30 minutes, followed by two washes in PBS plus 0.05% Tween (PBS-T). Cells were incubated with primary antibodies (α-Vimentin; 1:500; Invitrogen; PA1-16759) and 0.4 U/µL RiboLock RNase inhibitor (Thermo Fisher Scientific; EO0384) diluted in 2.5% BSA in PBS for 1 hour at room temperature. Cells were washed three times with PBS-T and incubated with conjugated secondary antibodies (α-chicken, AF647; Invitrogen; A21449) diluted 1:500 and 0.4 U/µL RiboLock RNase inhibitor in 2.5% BSA in PBS for 45 minutes at room temperature in the dark. The reverse transcription reaction was set up as follows: 1X RevertAid RT buffer (Thermo Fisher Scientific; EP0452), 250 µM dNTPs (New England Biolabs; N0447L), 1 µM biotinylated reverse transcription primer (oKC1015), 200 µg/mL recombinant albumin (rAlbumin) (New England Biolabs; B9200S), 0.8 U/µL RiboLock RNase inhibitor, and 4.8 U/µL RevertAid H minus Reverse Transcriptase (Thermo Fisher Scientific EP0452). The cells were incubated in the reverse transcription reaction overnight at 37°C shaking at 135 rpm. Cells were washed five times with PBS and incubated in 20 µg/mL Streptavidin (New England Biolabs; N7021S) and 100 µg/mL rAlbumin (New England Biolabs; B9200S) in PBS for 15 minutes. Cells were washed three times with PBS-T and fixed with 3% formaldehyde and 0.1% glutaraldehyde in 1X PBS for 30 minutes. Following three PBS-T washes, the cells were incubated in gap-fill and ligation solution at 37°C for 5 min, followed by 45 °C for 90 minutes. The gap-fill and ligation solution was set up as follows: 1X Ampligase buffer (Lucigen; A3210K), 200 µg/mL rAlbumin (New England Biolabs; B9200S), 0.1 µM padlock probe (oDF590), 50 nM dNTPs (New England Biolabs; N0447L), 0.02 U/µL TaqIT polymerase (Enzymatics; P7620L), 0.4 U/µL RNase H (Enzymatics; Y9220L), and 0.5 U/µL Ampligase (Lucigen; A3210K). Samples were then washed with PBS-T twice and incubated in rolling circle amplification (RCA) reaction solution overnight at 30°C. The RCA reaction solution was set up as follows: 1X Phi29 buffer (Thermo Fisher Scientific; EP0094), 250 µM dNTPs (New England Biolabs; N0447L), 200 µg/mL rAlbumin (New England Biolabs; B9200S), 5% glycerol, and 1 U/µL Phi29 DNA polymerase (Thermo Fisher Scientific; EP0094). Following RCA, cells were washed with PBS-T two times and again incubated with the same primary antibody solution minus RNase inhibitors for 45 minutes to enhance signal. Cells were again washed three times with PBS-T and incubated with the same secondary antibody solution for 45 minutes at room temperature in the dark. Following three PBS washes, wells were replaced with 2X SSC with 200 ng/mL DAPI and imaged for cellular phenotypes. Images were acquired using a Nikon Ti2 widefield microscope with 4 z-slices at 1.5 μm intervals. Following imaging, fluorophores were bleached using 1 mg/mL of lithium borohydride (LiBH_4_) (Radtke et al., 2020) for 1 hour. Cells were washed three times with PBS-T and incubated with primary antibody (α-integrin beta 1; 1:500; Abcam; ab30394) diluted in 2.5% BSA in PBS for 1 hour at room temperature. ITGB1-staining was nonspecific and thus only used for clustering purposes. Cells were washed three times with PBS-T and incubated with conjugated secondary antibodies diluted 1:500 in 2.5% BSA in PBS for 45 minutes at room temperature in the dark. Cells were again washed with PBS three times and imaged for cellular phenotypes in 2X SSC with 200 ng/mL DAPI. Of note, other markers were also included in both rounds of cellular phenotyping, but were not relevant to this study and are not reported here. Following imaging, fluorophores were again bleached in 1 mg/mL LiBH_4_ for 1 hour and then washed three times with 2X SSC.

Finally, *in situ* sequencing-by-synthesis was performed as described previously (Feldman et al., 2022). First, 1 µM sequencing primer (oDF663) in 2X SSC was hybridized for 30 minutes at 37°C, followed by two washes with PR2 (Illumina MS-103-1003). Cells were washed with PR2 and incubated in incorporation mix (MiSeq Nano kit v2 reagent 1, Illumina MS-103-1003) for 3 minutes at 60°C on a flat-top thermocycler. Cells were washed repeatedly with PR2, followed by three heated washes at 60°C for 5 minutes each. Prior to imaging of the first cycle of *in situ* sequencing, cells were incubated with FITC-conjugated anti-chicken secondary antibodies (diluted 1:500 in 2.5% BSA in PBS) for 45 minutes to label vimentin and aid in cellular segmentation. Cells were imaged in 2X SSC with 200 ng/mL DAPI. To proceed to the next cycle, samples were incubated in cleavage mix (Illumina kit reagent 4) for 6 minutes at 60°C. Samples were repeatedly washed with PR2, followed by two heated PR2 washes for 2 minutes at 60°C and one heated PR2 wash for 5 minutes at 60°C. Samples then returned to the incorporation step. Twelve cycles of *in situ* sequencing were performed. *In situ* sequencing images and cellular phenotypes were analyzed using Brieflow as previously described (Di Bernardo et al., 2025; Feldman et al., 2022; Feldman et al., 2019).

### PEG-mediated cell fusion

Cells were cultured to about 50-80% confluency. All solutions were prewarmed to 37°C. Cells were washed with PBS 2X and incubated in 50% PEG 8000 in PBS at room temperature for 2.5 - 3 minutes. The PEG solution was removed and cells were washed 3X with PBS and 2X with media. Cells were incubated at 37°C with 5% CO_2_ for 3 hours.

### Fibronectin coating

Glass-coverslips or 12-well glass-bottom plates (Cellvis; P12-1.5H-N) were incubated with 20 µg/mL fibronectin (Gibco; 33016015) in PBS for 1 hour at room temperature. Following incubations, the surfaces were gently washed with PBS and stored in PBS until ready to be used within the same day.

### Immunofluorescence and microscopy

For still-imaging of live cells, cells were seeded into 6-well glass-bottom plates (Cellvis; P06-1.5H-N) and induced for 72 hours. Hoechst was added to 1 µg/mL one hour before imaging. Images were acquired with a Nikon Eclipse microscope using a Plan Fluor 20X/0.5 NA objective.

For immunofluorescence experiments, cells were seeded onto uncoated or, if specified, fibronectin-coated glass coverslips. Cells were fixed with 4% formaldehyde in PBS for 10 minutes, washed with PBS, and permeabilized in PBS plus 0.2% Triton X-100 for 10 minutes. For ACA staining, cells were fixed instead with 4% formaldehyde in PHEM (60 mM PIPES, 25 mM HEPES, 10 mM EGTA, and 4 mM MgSO4). Coverslips were then blocked in AbDil (20 mM Tris-HCl, 150 mM NaCl, 0.1% Triton X-100, 3% bovine serum albumin, 0.1% NaN3, pH 7.5) for 30 minutes. Primary antibodies were diluted in AbDil and added to coverslips for 1 hour at room temperature or overnight at 4°C. Coverslips were washed 3 times with PBS. Cy2-, Cy5-, or Alexa647-conjugated secondary antibodies (Jackson ImmunoResearch Laboratories) were diluted 1:500 in AbDil along with 0.5 µg/mL Hoechst and applied to coverslips for 45 minutes at room temperature. For cells stained with phalloidin, Alexa Fluor™ 488 phalloidin (Invitrogen) in DMSO was diluted 1:5000 along with secondary antibodies. Coverslips were washed with PBS and mounted using ProLong^TM^ Gold antifade reagent (Invitrogen). Immunofluorescence images were taken on a DeltaVision Ultra (Cytiva) using a 40X/1.35NA or 60X/1.42NA objective. Images were analyzed using Fiji (ImageJ, NIH) and CellProfiler (Stirling et al., 2021).

The following primary antibodies were used: Human anti-centromere antibody (ACA, 1:200, Antibodies Inc, 15234), anti-alpha tubulin (1:1000; Abcam; ab52866), anti-phospho-myosin light chain 2 (ser19) (1:400, Cell Signaling; 3675S), anti-integrin beta 1 (1:250, Abcam; ab30394), anti-paxillin (1:200, Abcam; ab3127).

### Immunofluorescence analysis

For quantification of cortical and perinuclear actin intensity, preprocessing was first performed in Fiji, in which 7 in-focus sections were max projected and exported as tiffs. The tiffs were then imported into and quantified using CellProfiler (Stirling et al., 2021). The actin channel was background-subtracted by creating a mask of the cell-free areas of each image, calculating the median fluorescence intensity, and subtracting this value from the actin channel. The actin and microtubule channels were merged to create a combined image for automated cell segmentation. Major segmentation errors were manually corrected using the mem-mCherry channel as a guide. The nuclei were segmented using the DNA channel, again followed by manual correction. The cortex was defined as a 15-pixel-wide region inward from the cell boundary, the perinuclear region as a 7-pixel-wide region outward from the nuclear boundary, and the cytoplasm as the region between these two zones. For downstream analysis, the number of nuclei per cell was used to categorize cells as mononucleated or multinucleated, and cortical and perinuclear actin intensities were normalized to cytoplasmic actin intensity.

### Cell spreading assay

For spreading assays using inducible knockout cell lines, cells were seeded in 6-well plates and induced for 72 hours. For assays with PEG-fused cells and unfused control cells, cells were seeded in 6-well plates, cultured for 2 days, and fused approximately 3 hours prior to the assay. For the spreading assay, cells were trypsinized, resuspended in media, and counted using a Z2 Coulter Counter (Beckman Coulter). Cells were spun down and resuspended to equal densities in CO_2_-independent media (Gibco) supplemented with 10% FBS, 100 U/mL penicillin and streptomycin, and 2 mM L-glutamine and plated onto fibronectin-coated wells in a glass-bottom 12-well plate (Cellvis; P12-1.5H-N). Cells were imaged immediately at 37°C on a Nikon Eclipse microscope using a Plan Fluor 20X/0.5 NA objective. Phase-contrast images were acquired in 5-minute intervals for 1 hour, followed by 20-minute intervals for 23 hours. Image brightness and contrast were adjusted in Fiji (ImageJ, NIH). Cells were segmented using Cellpose-SAM (Pachitariu et al., 2025) and processed using CellProfiler (Stirling et al., 2021) to measure cell form factor. Of note, segmentation of multinucleated cells was occasionally inaccurate, limiting reliable quantification. Representative segmentation masks are therefore presented to illustrate performance for these conditions.

All spreading assays were performed in parallel experiments, and control data are reused across figures for visualization purposes. Three independent replicate experiments were performed; however, data shown in supplemental figures were included in only two of the three replicates, and accordingly, only the control data from those two replicates are displayed.

### Nuclear tracking analyses

Analyses of nuclear motility were performed on the brightfield images acquired during spreading assays. Nuclei were tracked in Fiji using the MTrackJ plugin (Meijering et al., 2012). Tracking was initiated 35 minutes following the start of acquisition, ensuring cells had settled and attached, and continued until the end of the movie (1445 min). Cells that died before the end of the movie were thus excluded from this analysis. In the case where a cell divides, one of the daughter nuclei was selected and tracked for the remaining frames. Displacement of the nuclei reflects both movement of the nuclei within the confines of the cell body as well as cellular movement. For conditions that produce multinucleated cells, only multinucleated cells were quantified.

### Quantitative mass spectrometry

For whole-cell quantitative proteomics, control and KCTD10 or ECT2 iKO cells were induced for 96 or 72 hours, respectively. For KCTD10 and ECT2 iKOs, cells were harvested by mechanical shake-off to isolate weakly-adherent cells, thereby enriching for cells with successful gene knockout. To prevent simultaneous enrichment of mitotic cells, cells were arrested in S phase by blocking with 2 mM thymidine for 24 hours prior to harvesting. Controls were also subjected to thymidine block, but were harvested instead via PBS-EDTA-mediated detachment. Cell pellets were washed once with PBS and once with lysis buffer (50 mM HEPES pH 7.4, 1 mM EGTA, 1 mM MgCl_2_, 300 mM KCl, 10% glycerol), snap-frozen in liquid nitrogen, and stored at - 80°C. Three biological replicates were collected for each condition.

To prepare protein extracts, cells were thawed on ice in lysis buffer supplemented with 0.05% NP-40, 20 mM beta-glycerophosphate, 5 mM sodium fluoride, 0.4 mM sodium orthovanadate, 1X cOmplete protease inhibitor cocktail (Roche), and 1 mM phenylmethylsulfonyl fluoride (PMSF). Cells were lysed via sonication and clarified by centrifugation at 14,800 rpm for 30 minutes at 4°C. Trichloroacetic acid (TCA) was added to the supernatant to a final concentration of 20% and incubated on ice overnight to precipitate proteins. Samples were then centrifuged at 14,800 rpm for 30 minutes at 4°C, washed two times with cold acetone, and dried using a speedvac.

To prepare samples for mass spectrometry, pellets were resuspended in 5% SDS, 50 mM TEAB, pH 8.5, 20 mM DTT and incubated at 95°C for 10 min. After samples cooled back down to room temperature, iodoacetamide was added to 40 mM and the samples were incubated in the dark at room temperature for 30 minutes. Phosphoric acid was then added to 2.5% to quench the alkylating reaction. Six volumes of S-trap binding buffer (90% MeOH, 100 mM TEAB, pH 7.55) were added and then loaded onto S-trap mini columns (Protifi) and spun at 4,000 x g for 30 seconds. Samples were washed with S-trap binding buffer 4 times. Following the final wash, columns were dried by spinning at 4,000 x g for 1 minute. Proteins on the column were digested with 1 µg trypsin in 20 µL 50 mM TEAB pH 8.5 overnight at 37°C in a humidified chamber. The peptides were eluted using 40 µL 50 mM TEAB pH 8.5, followed by 40 µL 0.2% formic acid, and then 40 µL 50% acetonitrile. The eluted peptides were pooled, quantified using a Quantitative Fluorometric Peptide Assay kit (Pierce), and lyophilized.

For quantitative mass spectrometry, peptides were labeled using the TMT10plex labels. Peptides were dissolved in 50 mM TEAB pH 8.5 and labeled at a 10:1 label:peptide w/w ratio in 30% acetonitrile for 1 hour at room temperature. The labeling reaction was quenched with hydroxylamine for 15 minutes at room temperature. The samples were then pooled together and lyophilized. Next, pooled TMT-labeled peptides were fractionated using the Pierce High pH Reversed-Phase Peptide Fractionation kit (Pierce) according to the manufacturer’s instructions and lyophilized. Fractions were resuspended in 0.2% formic acid and loaded onto an Exploris 480 Orbitrap mass spectrometer. Proteome Discoverer 2.4 (Thermo Fisher Scientific) was used for protein identification and TMT quantification. The mass spectrometry proteomics data have been deposited to the ProteomeXchange Consortium via the PRIDE (Perez-Riverol et al., 2025) partner repository with the dataset identifier PXD080810 and https://doi.org/10.6019/PXD080810 (ECT2) and PXD080816 and https://doi.org/10.6019/PXD080816 (KCTD10).

